# Rescue of bacterial motility using two and three-species FliC chimeras

**DOI:** 10.1101/2024.12.02.626473

**Authors:** Jacob Scadden, Pietro Ridone, Divyangi Pandit, Yoshiyuki Sowa, Matthew AB Baker

**Affiliations:** School of Biotechnology and Biomolecular Sciences, University of New South Wales, Sydney, Australia; Department of Frontier Bioscience, Hosei University, Tokyo, Japan; Research Center for Micro-Nano Technology, Hosei University, Tokyo, Japan

## Abstract

The bacterial flagellar filament acts as a propeller to drive most bacterial swimming. The filament is made of flagellin, known as FliC in *Escherichia coli*. FliC consists of four domains, the highly conserved core D0 and D1 domains and the hypervariable outer D2 and D3 domains. The size and structure of the outer domains varies, being completely absent in some bacterial species. Here, we sought to identify outer domains from various species that are compatible with the ability of *E. coli* K-12 FliC to form filaments capable of supporting motility. We calculated a phylogeny of 210 representative flagellin amino acid sequences and generated a series of FliC variants, including outer-domain-deleted forms and eleven chimeric FliC mutants using domains from *E. coli K-12, Salmonella* Typhimurium, *Pseudomonas aeruginosa, Collimonas fungivorans, Helicobacter mustelae* and *Mesorhizobium* sp. ORS3359 in various combinations. Notably, two of the chimeric *fliC* mutants rescued motility in a *fliC*-disrupted *E. coli* K-12 strain, both of which contained the *S*. Typhimurium D2 domain. Overall, we demonstrate that, while most FliC chimeras did not support motility, interchangeability of the outer domains can produce filaments that provide motility, providing insights to guide the design of synthetic flagellins.

**Importance:** Flagellin is a key protein forming the filament of the bacterial flagellar motor which powers most bacterial swimming. Flagellin can have hypervariable domains which can alter motility in different environments and provide immune evasion. Here we engineered two flagellin chimeras that could drive motility. This indicates that the flagellin outer domains can be exchanged, to some degree, allowing us to refine rational design approaches for engineering of bacterial swimming. Our work shows the challenges to overcome when combining flagellins from different species and provides evidence that domain switched flagellins can form filaments.

## Introduction

Motility provides organisms the ability to exploit new niches, to evade predation and to find increased opportunities for the exchange of genetic material (1). The flagellar filament is a major component of the bacterial flagellar motor (BFM) taking the form of the bacterial flagellum that consists of a long helical protein filament which is formed from approximately 30,000 flagellin subunits, known as FliC in *Escherichia coli* K-12 (2,3). FliC interacts with both the flagellar hook junction proteins (FlgK and FlgL) and filament cap protein (FliD) (4–6).

FliC from *E. coli* K-12 is comprised of four domains, D0, D1, D2 and D3 (7,8). Domains D0 and D1 are found on the N- and C-termini of the protein and are highly conserved across multiple bacterial phyla, such as *Proteobacteria, Bacillota* and *Actinobacteria* (9). These “core” domains are formed of α-helices and interact with each other as part of the supercoiled filament (7,8,10,11). Within the D0 domain is a 22-residue segment which is the recognition signal for the flagellar export chaperone, FliS (12). Domains D2 and D3 in *E. coli* K-12, also termed “outer domains”, are variable in both sequence and length even at a species level (3). These outer domains consist primarily of β-sheets and were shown in *Salmonella* Typhimurium to have no interactions between each other (7,13). In some cases, flagellin monomers from other bacterial species such as *Bacillus cereus, Agrobacterium tumefaciens* and *Treponema pallidum* do not contain a D2 or D3 domain but still produce a functional flagellar filament capable of motility (9,14–18). It has been shown that within the genera *Escherichia* and *Salmonella* there is a large level of diversity in the outer domains, suggesting that the outer domains are acquired via the process of evolution (19,20). This implies that the outer domains of flagellin from diverse species can be incorporated into host flagellins to provide competitive advantages that increase efficiency of motility in different environments or that allow evasion of recognition by the immune system.

In *Pseudomonas aeruginosa* PAO1 the outer domains of flagellin increase the motility of bacteria in more viscous environments due to inter-domain interactions (21). Likewise, the outer domains *E. coli* O157:H7, *E. coli* O127:H6, *Achromobacter* (which have an additional “D4” domain) and *Sinorhizobium meliloti* (which has only a D2 outer domain) can dimerise or tetramerise to form a screw-like surface which increases motility through stabilisation an intermediate waveform and prolonging tumbling (22). Furthermore, interactions of the outer domains of *E. coli* can increase tumbling time (22). Inter-species flagellin chimeras between *Salmonella* strains (serovar Typhimurium strain 14028s and serovar Enteritidis strain P125109) were able to form functional filaments (23). Of 13 FliC variants, four were motile, and it was shown that alternating the outer domains could impact motility (23).

One of the main research focuses on the flagellin outer domains is their use in immune activation and protein display. *S*. Typhimurium FliC is capable of activating the Toll-Like Receptor 5 (TLR5) (24). In a comprehensive review by Hajam *et al*., the authors identified 29 studies where flagellin was used as an immune adjuvant for bacterial, viral, parasitic and other miscellaneous antigens (25). This use of recombinant flagellin as a surface-display tool for immune activation has led to a number of vaccines being developed and used in human clinical trials (26,27). However, although chimeric FliC has been used in immunological studies, the effect of manipulating the outer domains on bacterial motility is not well understood.

Here, we tested the ability of chimeric FliC to support motility where we replace the outer domains of *E. coli* K-12 with outer domains from flagellins of a diverse set of bacterial. We show that the outer domains of *E. coli* K-12 FliC are not required to drive swimming motility and that two forms chimeric FliC can form functional filaments that support swimming motility.

## Materials and Methods

### Bacterial strains, plasmids and growth conditions

Table S1 describes the strains and plasmids used in this study. The *E. coli* strain SYC29 was constructed from the chemotactic wild-type strains RP437, using the λ RED recombination system, as previously described (28,29). *E. coli* strains were grown in Lysogeny Broth (LB) (10 g/L NaCl, 10 g/L Bacto-tryptone, 5 g/L Yeast Extract). For 2% (w/v),0.3%, 0.35% and0.4% (w/v) LB agar plates 20 g/L, 3 g/L, 3.5 g/L or 4 g/L agar was added, respectively. *E. coli* strains were grown in Tryptone Broth (TB) to aid in induction of motility (5 g/L NaCl, 10 g/L Bacto-tryptone). Antibiotic selection used ampicillin (AMP) (50 µg/ml), chloramphenicol (CAM) (25 µg/ml) and/or tetracycline (TET) (10 µg/ml). Induction for gene expression from plasmids required L-arabinose (0.02% (w/v)) and/or Isopropyl ß-D-1-thiogalactopyranoside (IPTG) (0.1 mM).

### Phylogenetic analysis of FliC

To investigate the structural diversity of the outer domains of flagellins and their presence/absence across the tree of life, eight flagellin amino acid sequences (Table S2) were used as reference sequences for a BLASTP search in Uniprot (30) (E-Threshold - 10; Matrix – BLOSUM62; Filter – None; Gapped – Yes; Hits – 1000; HSPs per hit – All). All hits for each BLASTP search were selected and used in ID mapping from database UniProtKB AC/ID to UniRef90 (31). The clusters were sorted by member size and each list was manually curated to select 210 representative flagellin sequences for further phylogenetic analysis, in addition to 14 FliD amino acid sequences as an outgroup. The 224 sequences were downloaded in FASTA format and uploaded to CIPRES Science Gateway (32). Alignment of the representative sequences was performed using MUSCLE (3.7) (33) and viewed using AliView (34). The alignment was used to generate a phylogenetic tree using IQTree (2.3.2) (35) using CIPRES (for parameters see Table S3) (32). 682 representative flagellins were downloaded from the supplementary information from Fields *et al*. (20), 605 were available on the Uniprot database and the amino acid sequences used in a MUSCLE alignment (33). A FastTree2 (36) phylogenetic tree was generated using CIPRES (32). Phylogenetic trees were visualised and annotated in ItoL (v6) (37).

### Design and structural prediction of chimeric FliC

To study the effect of outer domain manipulation on bacterial motility, a rational design approach for each chimeric FliC candidate was taken. It was decided to maintain the D0 and D1 domains of *E. coli* K-12 FliC (UniProt - P04949) (EEEE) in order to preserve the export recognition sequence found in the D0 domain (12). Therefore, to understand if these outer domains are functionally required for flagellum formation, the first *E. coli* K-12 *fliC* variants designed were the *fliC_*Δ*D3* (EEE) and *fliC_*Δ*D2/3 (EE)*, with deletions of residues 195-300 and 174-405, respectively. For the two-species and three-species FliC chimeras, five flagellin amino acid sequences were used from across the bacterial tree of life, *Helicobacter mustelae* (P50612), *Mesorhizobium* sp. ORS3359 (A0A090GIX0), *Pseudomonas aeruginosa* (P21184), *Collimonas fungivorans* strain Ter331 (G0AIL6) and *Salmonella* Typhimurium strain LT2 (P06179). The D2 and/or D3 domains of these flagellins were determined manually and used to replace the D2 and D3 domains of *E. coli* FliC. In the case of *S*. Typhimurium additional chimeras were made using the D2 domain of *S*. Typhimurium and the D2 domains (in place of the D3 domains) of *Mesorhizobium* sp. ORS3359, *P. aeruginosa, C. fungivorans* strain Ter331 were used. Structural prediction for all domain deleted and chimeric FliC sequences was performed using AlphaFold3 (v3.0.1) (38).

### Cloning of FliC chimeras into expression vector

Reverse translation of each FliC chimera from amino acid to nucleotide codon optimised sequence for *E. coli* was performed in Benchling (Benchling Inc). Gene fragments for all chimeras and outer domain deleted FliC were order using TWIST (Decode Bioscience) with EcoRI and NotI restriction sites incorporated in the 5’ and 3’ regions, respectively. Digestion of gene fragments and pET21(+) (Merck) expression vector was performed using EcoRI-HF and NotI-HF (NEB), followed by Antarctic phosphatase (NEB) treatment of pET21(+), as per the manufacturer’s instructions. Ligation was carried out using T4 ligase (NEB) as per manufacturer’s instructions and incubated at room temperature overnight. After heat inactivation, the ligation mix was transformed into to chemically competent *E. coli* 10-β (NEB) as per manufacturer’s instructions. Transformations were incubated at 37°C for 1 hr and plates on LB plates with AMP. Colonies were screened using pET21_F and pET21_R primers and Q5 Polymerase (NEB) (See Table S4 for primer details and PCR conditions). Positive colonies were grown overnight in LB medium with AMP and plasmids were purified using a Qiagen Miniprep Kit, following manufacturer’s protocol. Purified plasmid was sent for Sanger sequencing performed by the Ramaciotti Centre for Genomics (UNSW, Sydney). The confirmed plasmids were then transformed into chemically competent *E. coli* SYC29 pDB108 (Δ*motAB, fliC*::*tetR*) and screened on LB plates containing AMP, CAM and TET. This strain was selected due to its inducible motility characteristics. Positive colonies were screened using the same PCR method as previously described and grown overnight in LB medium containing AMP, CAM and TET.

### Assessing the motility of chimeric FliC expressing *E. coli*

Swim plate (0.30%, 0.35% and 0.40% agar) motility assays were performed with required antibiotics and inducers as previously described (39). Plates were inoculated with 1 µl induced overnight culture and incubated at 30°C for 16 hrs. Plates were imaged using a Bio-rad Gel Doc XR with Gel documentation system with analysis of Image Lab 6.0.1 (No filter, White Epi Illumination). Swim ring diameters were measured using ImageJ (FIJI) (40). Data were analysed and displayed using GraphPad 2.0 (version 9.5.1).

### Free swimming assay for motile chimeric FliC expressing *E. coli*

Mutant FliC expressing *E. coli* were grown overnight in LB medium containing appropriate antibiotics and inducers at 30°C. A 1% inoculum was grown in TB medium until OD_600_ 0.5. A tunnel slide was formed using tape and a glass cover slip and 30 µl of sample was flown through, as previously described (41). Cells were visualised using a phase contract microscope (Leica) and imaged using a Nikon camera. For each sample 10 videos of 400 frames (20 frames/second) were recorded. Recordings were analysed in a custom programme in LabVIEW (National Instruments) (39) to quantify the swim speeds of each motile construct. GraphPad 2.0 (version 9.5.1) was used to analyse and display data.

### Flagella staining of chimeric FliC expressing *E. coli*

Staining of flagellar filaments was performed using Remel ^™^ Flagella stain (Thermofisher) as per the manufacturers protocol with minor changes. Strains of interest were inoculated on 0.3% LB agar plates, with appropriate antimicrobials and inducers, and incubated for 16 hrs at 30°C. To the edge of each swim ring 30 µl of LB medium was added and incubated for two mins at room temperature after which 10 µl was spotted on a glass slide. After this step the protocol was followed exactly. Samples were visualised using a Leica DM750 microscope at 100x magnification and images captured using a MEE-500 camera (Yegren, China) and the camera app on Windows 10 (v 10.0.1.19045). Filament lengths were measured using ImageJ (FIJI) (40) using the freehand line tool. Data were analysed and displayed using GraphPad 2.0 (version 9.5.1).

## Results

### Flagellins cluster according to primarily phyla

To study the structural diversity of flagellin outer domains, 210 representative flagellin amino acid sequences from across the bacterial tree of life were selected, aligned and used in the generation of a phylogenetic tree (Figure 1). Of the representative flagellin sequences selected 53% (112/210) contained the outer D2 and D3 domains. The majority of the flagellins clustered according to the phyla or class to which the bacteria encoding the respective flagellin gene belonged, also seen in phylogenetic analysis of 605 flagellin sequences from Fields *et al*., (20) (Figure S1A and B). In *Spirochaetota* (12/12) and *Actinomycelotota* (10/10), all flagellins did not contain the outer domains (Figure 1, red branches). In *Bacillota*, none of the flagellins contained outer domains, with one exception (53/54). The classes *Epsilonproteobacteria* (20/20) and *Beta/Gammaproteobacteria* (66/67) all clustered within their respective class and all contained the outer domains, with one exception (Figure 1, blue branches). *Deltaproteobacteria* and the *Fibrobacteres-Chlorobi-Bacteroidetes* (FCB) superphylum had flagellin sequences both with and without outer domains and the clading of these was split on the presence or absence of the outer domains. *Alphaproteobacteria* flagellins, in contrast, were the only group where sequences with and without the outer domains where mixed in the one clade (10 with and 12 without, respectively). In *Deltaproteobacteria* and the FCB superphylum, 64% (9/14) and 42% (5/12), respectively, contained outer domains.

**Figure 1.**
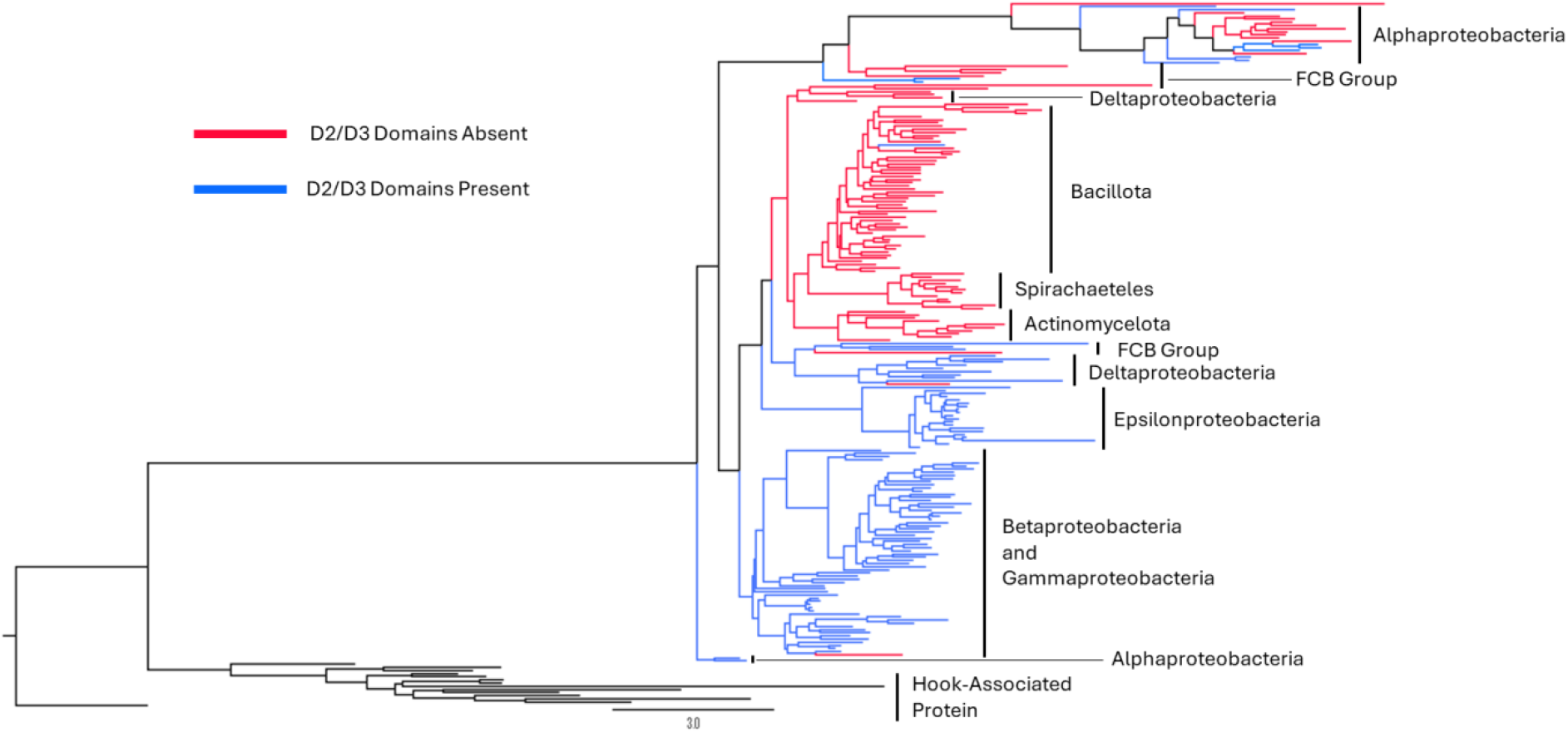
Phylogeny of 210 representative flagellin sequences, with 14 flagellar hook associated proteins as an outgroup. Branches are coloured based on the presence (Blue) or absence (red) of the D2 and D3 domains.

The majority of flagellins containing outer domains were isolated from animal and water associated environments (Figure S2A), whereas flagellins lacking the outer domains were found across soil, animal associated and aquatic environments (Figure S2B). This was also observed when comparing isolation sources, with a higher percentage of flagellins that contained outer domains were isolated from animal and plant sources (63% and 55%, respectively) (Figure S2C). However, flagellins lacking the outer domains were found predominantly in soil isolates (70%). There was a near equal percentage of flagellins containing outer domains and flagellins with no outer domains isolated from aquatic environments (51% and 49%, respectively). Within each phylum-based cluster there was a minimal association of a flagellin type with a particular environment, with the exception of *Epsilonproteobactera*, 88% of which were isolated from animal sources, and *Alphaproteobacteria*, which were all either from aquatic or plant associated environments.

### Outer domain deleted and chimeric flagellin can rescue motility in *E. coli* K-12

First, to investigate the potential redundancy of the outer domains for bacterial motility we deleted the nucleotide sequence coding for the outer domains from *E. coli* K-12 *fliC*. The nomenclature used for all flagellin outer domain deletions and chimeric variants created in this work represents each domain (D0-D3) as a single character of the first letter of the genus to which the flagellin is derived, thus wild type *E. coli* K-12 flagellin is termed EEEE. Structural prediction of these outer domain deletions indicated that sequential removal of these domains, removing D3 to generate EEE and both D2 and D3 to generate EE, would not effect the structure of the core domains (Figure 2A, Figure S3). All PAE scores indicated high confidence at the N- and C-termini with reduced confidence in the central section of the protein, correlating to the position of the outer domains (Figure S4A-N). We then designed chimeric *fliC* variants using the D0 and D1 domains of *E. coli* K-12 and the outer domains of *S*. Typhimurium (EESS/EEES), *H. mustelae* (EEH/EEEH) and *C. fungivorans* (EEC/EEEC) (Figure S3A and D). The D0 and D1 domains were maintained as they are in *E. coli* K-12 to retain the export recognition site found between residues 26-47, given our use of *E. coli* K-12 proteins for all other elements of the motor, including flagellin export (12). *S*. Typhimurium (*Gammaproteobacteria*) (EESS/EEES) and *C. fungivorans* (*Betaproteobacteria*) (EEC/EEEC) were selected as members of the *Pseudomonadota* phylum, which clustered in the same clade as *E. coli* K-12 FliC (Figure 1). The outer domains of *H. mustelae* (EEHH/EEEH), belonging to *Epsilonproteobacteria*, were selected due to being in a cluster where all members contained the outer domains (Figure 1). The structures of these chimeric FliC variants were predicted using AlphaFold3 and these structural predictions showed that the tertiary structure of the core domains were conserved (Figure 2B, Figure S3).

**Figure 2.**
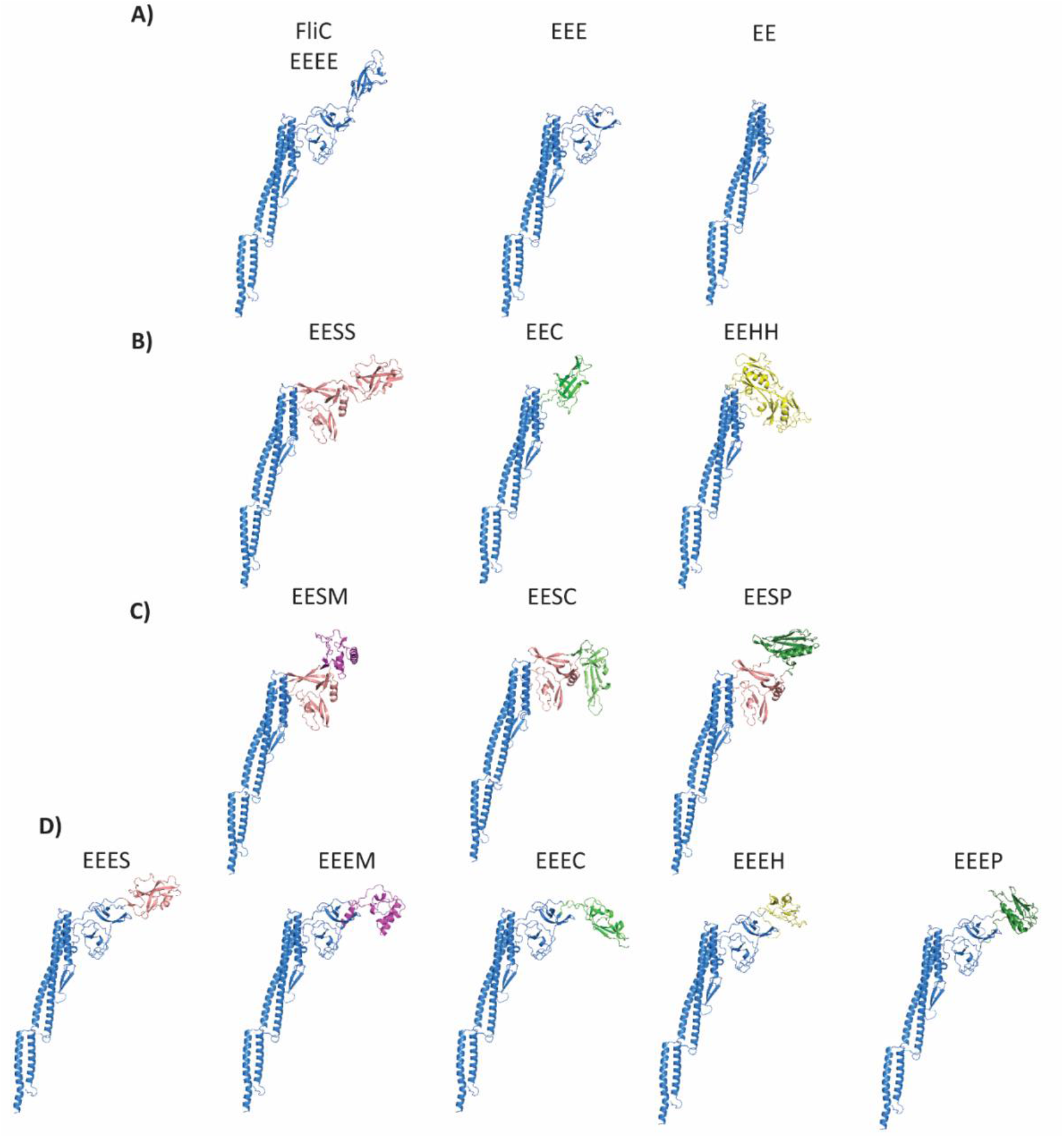
AlphaFold3 predictions of chimeric and outer-domain-deleted *E. coli* K-12 FliC monomers designed for this study. A) Predicted structures of wild type, D3-deleted and D2/3-deleted flagellin. B) Predicted structure of *E. coli* K-12 D0 and D1 domains and single species outer domains. C) Predicted structures of *E. coli* K-12 D0 and D1 domains and multi-species outer domains.D) Predicted structures of E. coli K-12 D0, D1 and D2 domains and single species outer domains. D) Predicted structures of *E. coli* K-12 D0, D1 and D2 domains with the D3 domain replaced with the outmost domain of a single species flagellin. *E. coli* K-12 (Blue), *H. mustelae* (Yellow), *C. fungivorans* (Light Green), *S*. Typhimurium (Pink), *Mesorhizobium* sp. OR3359 (Purple) and *P. aeruginosa* (Dark Green).

In addition to direct replacement of the outer domain replacement with a domain from a single species, we decided to engineer three-species FliC chimeras. This was to assess the inter-compatibility of the outermost D3 domain. The D2 domain of *S*. Typhimurium was used for all D2 domains in these three-species FliC constructs (Figure S3). The D3 domain was replaced with the outer domain from *C. fungivorans* strain Ter331 (EESC), *P. aeruginosa* (EESP), *Mesorhizobium* sp. ORS 3359 (EESM) (Figure 2C). The D3 domain from *C. fungivorans* strain Ter331 was chosen due to its FliC phylogenetic clading with the FliC from *E. coli*. Alternately, the D3 domain from *P. aeruginosa* and *Mesorhizobium* sp. ORS 3359 were chosen as previous work showed that the outer domains of *P. aeruginosa* PA01 and *Sinorhizobium meliloti* formed ridged filaments formed by interaction between the flagellin outer domains (Figure 2C) (21,22). Overall, chimeric and outer-domain-deleted FliC ranged in size from 266 to 535 residues in length, with the outer D2 and D3 domains ranging from 107 to 269 amino acids in length (Table S4). Additionally, the outermost D3 domain of the *E. coli* K-12 flagellin was replaced with the outermost domain from *S*. Typhimurium, *Mesorhizobium* sp. ORS 3359, *C. fungivorans, P. aeruginosa* and *H. mustelae* (Figure 2D).

Next, we investigated whether chimeric FliC could form functional filaments and therefore rescue motility in filament-disrupted strains (*fliC::tetRA*). Plasmids expressing FliC mutants or flagellar stators were co-transformed into an *E. coli* Δ*motAB fliC::tetRA* strain. Motility of outer-domain-deleted and chimeric FliC expressing strains was confirmed using swim plate motility assays. Both outer domain deletions produced swim rings (Figure 3). The average swim ring diameter when both outer domains were deleted (EE) was larger, 14.5 ± 2.5 mm, than that with only the D3 domain deleted, 11.9 ± 2.5 mm, however this difference was not statistically significant (Figure 3). However, both these outer-domain-deleted flagellin mutants produced significantly smaller swim rings than wild type FliC. Likewise, motility of the two-species outer domain FliC mutants, EESS, EEH and EEC, was confirmed using a swim plate motility assay. Only EESS was motile, with a swim ring diameter of 9.9 ± 2.1 mm (Figure 3). EEC and EEH produced no swim rings (Figure 3). Only one of the three-species chimeras were observed to be motile. The FliC chimera that contained the D3 domain from *Mesorhizobium* sp. OR3359 consistently produced a swim ring diameter of 10.6 ± 0.3 mm, whereas EESC and EESP were not motile (Figure 3). All flagellin chimeras with the D3 domain of *E. coli* K-12 replaced by the outermost domain of another bacterial flagellin were non-motile.

**Figure 3.**
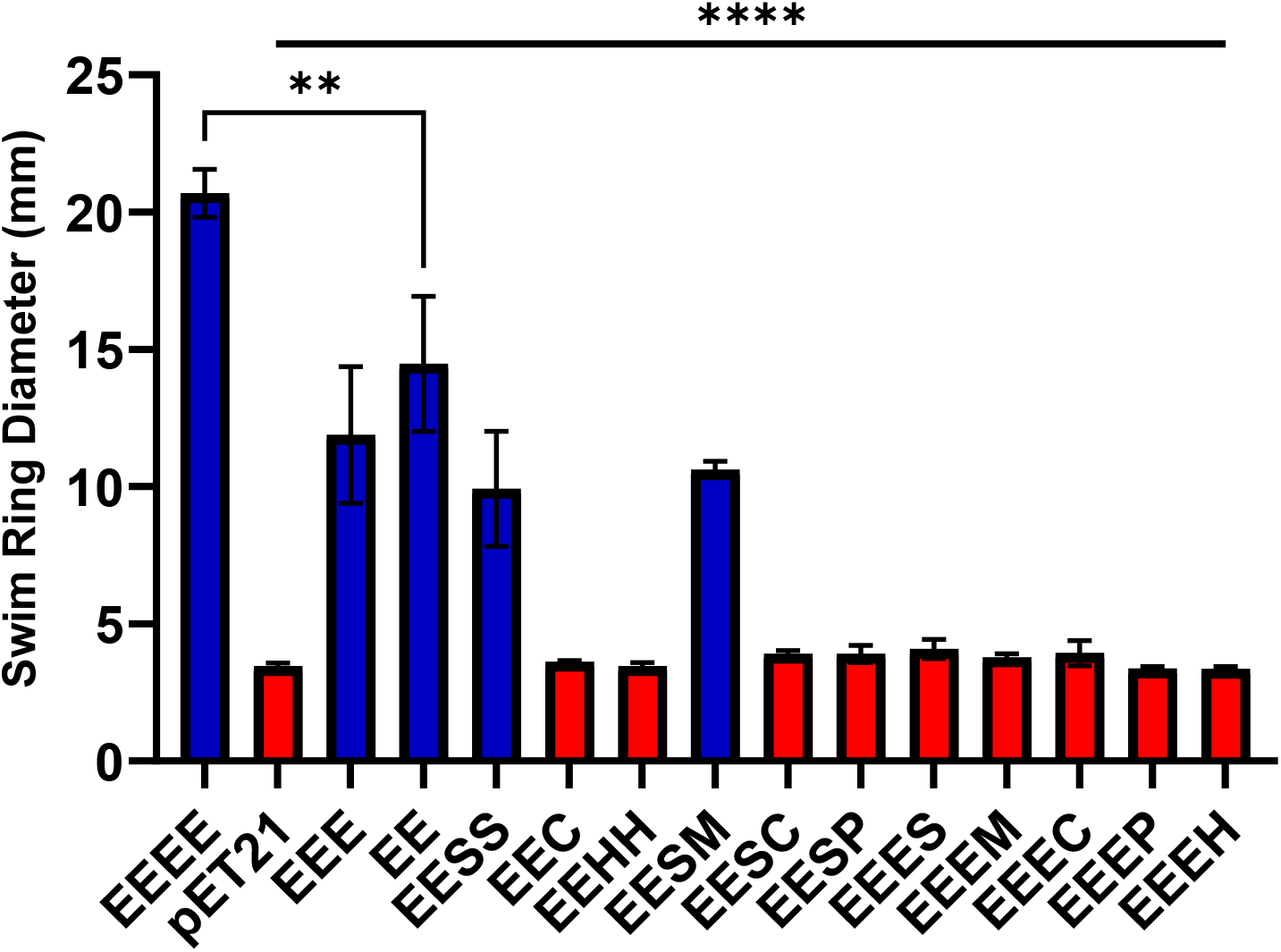
Mean swim ring diameter of cells expressing each chimeric and outer-domain-deleted FliC recorded from three independent 0.3% swim plates incubated at 30°C for 16 hrs. Motile and non-motile flagellin variants expressing chimeric FliC are indicated by blue and red bars, respectively. Error bars indicate standard error of the mean. A two-way ANOVA comparing each column to the FliC value was used to generate the p values (^**^ = ≤ 0.01, ^***^ = ≤ 0.001).

As the viscosity of agar plates was increased, the swim ring diameter decreased significantly (Figure S5A), with the overall relationship between the chimeric FliC expressing strains and WT FliC remaining the same (Figure S5 B). However, when normalised to the swim ring diameters using 0.3% agar there were no significant differences in the percentage of the swim ring diameter between any motile flagellin variant and wild type at either 0.35% or 0.4% agar (Figure S5B).

### Motile FliC chimeras that produced significantly lower swim speeds

Free swimming assays were used to measure the swimming speeds for each of the motile FliC strains that were generated as part of this work. Strains expressing EEEE (11.2 ± 2.3 µm/s) had swim speeds significantly faster than the other motile variants (p value < 0.0001) (Figure 4). Strains expressing EEE (2.4 ± 1.7 µm/s), EE (1.9 ± 0.6 µm/s), EESS (1.7 ± 0.8 µm/s) and EESM (3.44 ± 1.5 µm/s) had significantly lower swim speeds.

**Figure 4.**
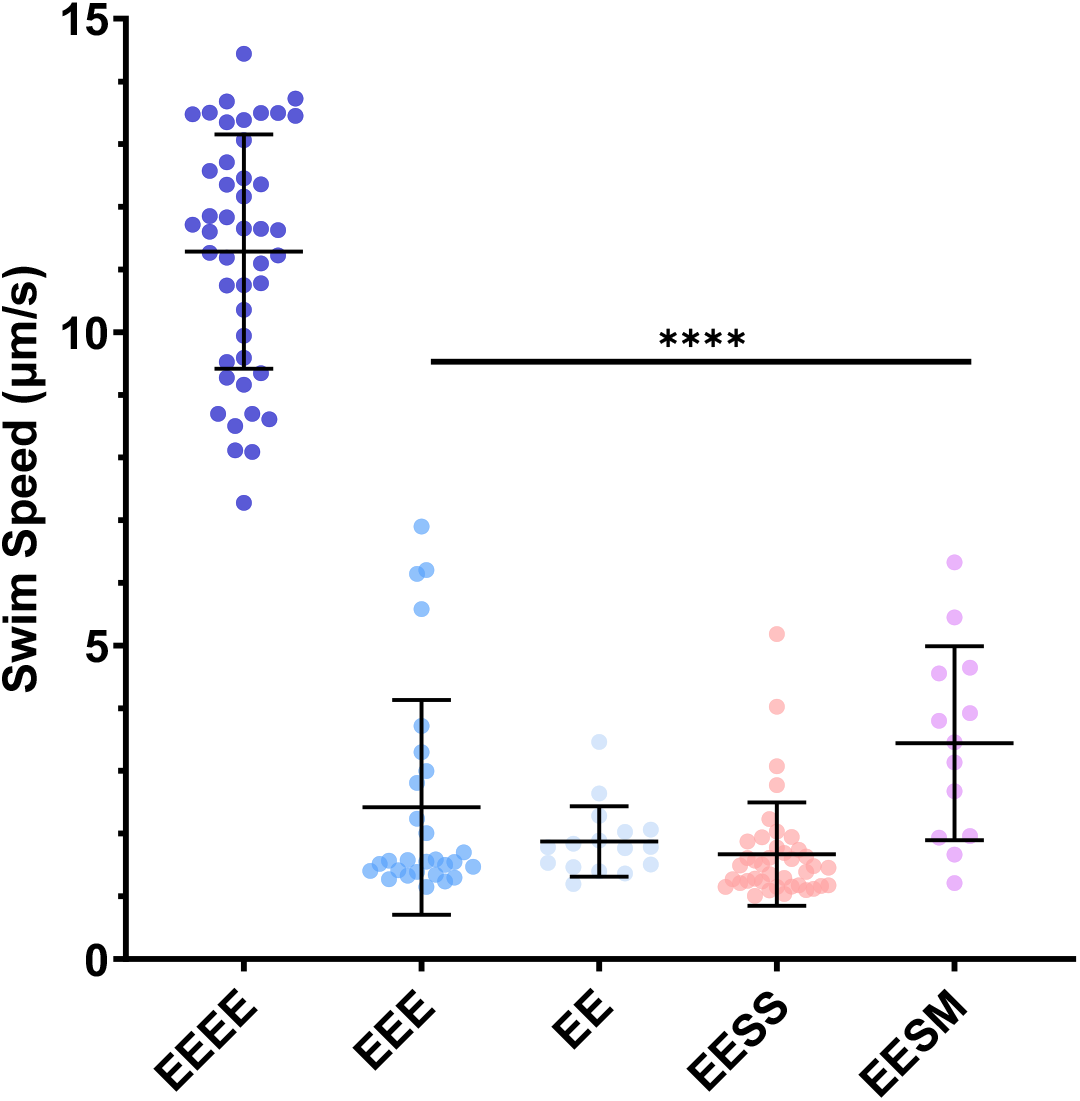
Free-swimming speeds for motile chimeric and truncated flagellin in µm/s. Error bars indicate standard deviation. A two-way ANOVA comparing each column to EEEE was used to generate the p values. Cells expressing wild-type FliC (EEEE) are significantly different to all other FliC variants(p value ≤ 0.05) (^****^ = ≤ 0.0001).

### Crystal violet staining of filaments shows formation of chimeric filaments

Staining of flagella confirmed export of flagellin and the formation of filaments. Filaments were observed in all motile strains (Figure 5A). For all non-motile strains there were no observable filaments. Filament length was determined for all motile strains with EEEE (FliC), producing the longest filaments at 5.7 ± 1.6 µm (Figure 5B). The outer domain deleted contructs EEE and EE produced significantly shorter filaments at 2.7 ± 0.9 µm and 1.9 ± 0.5 µm, respectively. Shorter filaments were also formed in the two-species chimera EESS (2.8 ± 1.1 µm) and in the three-species chimera EESM (3.4 ± 1.1 µm) (Figure 5B). All motile cells expressing domain-deleted or chimeric FliC had significantly shorter filaments than cells expressing wild-type FliC (EEEE).

**Figure 5.**
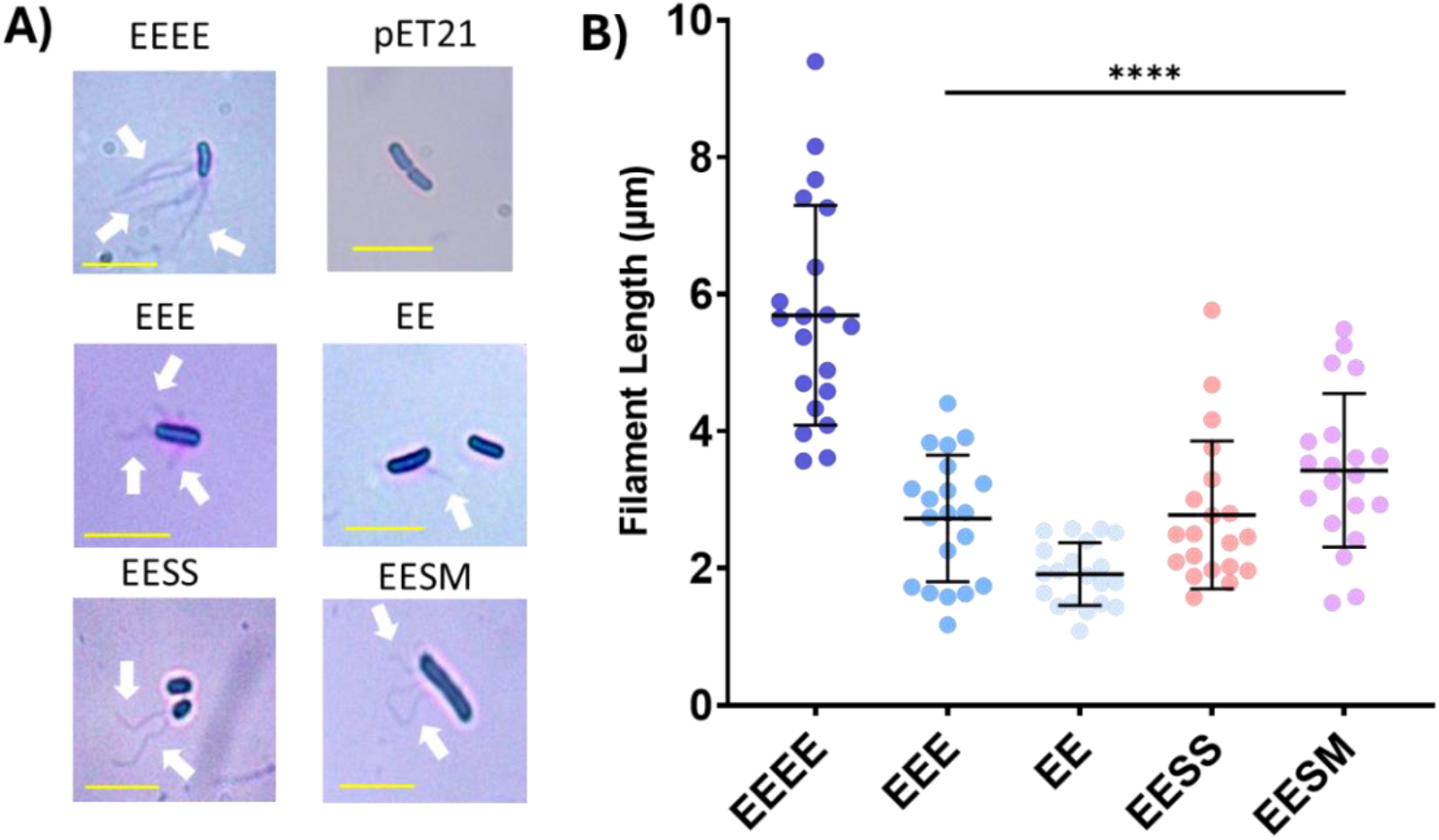
A) Analysis of filament export and assembly. Reprentative images of motile strains after flagella staining. The pET21 empty vector is shown as a representative non-motile variant. Scale bar (Yellow) represents 5 µm. White arrows indicate presence of flagella. B) Lengths of FliC chimeric filaments. Error bars represent standard deviation. A two-way ANOVA comparing each column to FliC was used to generate the p values (^****^ = ≤0.0001).

## Discussion

In this study, we investigated whether expression of *E. coli* FliC with deleted or replaced outer domains (D2 and D3) could rescue motility in *fliC*-deleted strains of *E. coli*. We observed that flagellin outer domains were not required for motility in *E. coli* and that two and three-species FliC outer domain chimeras are able to form functional filaments. From our phylogenetic analysis we showed that bacteria have evolved flagellar filaments with and without outer domains. Th presence or absence of these domains is distributed in a way that correlates with bacterial clades. A greater proportion of flagellins with outer domains were isolated from animal sources. Previous work has shown that the presence of these hypervariable domains can aid in immune evasion (42). Horizontal gene transfer of complete sets of flagellar genes has been demonstrated as well as evidence of specific recombination of the flagellin outer domains (43,44,19). Here, we we constructed FliC with chimeric or deleted outer domains to show that recombination of outer domains can form functional filaments, thus providing evidence that recombination events can occur across multiple genera with compatible outer domains.

### FliC outer domains can affect bacterial motility

Although originally thought that the outer domains have no interaction with the core domains or each other (7,13), recent studies have shown that in some cases there are inter-domain interactions which help to stabilise the filament to aid motility (21). In Kreutzberger *et al*., this phenomenon was identified in *Achromobacter* filaments where the outer domains exist in four conformations and can form a tetrameric sheath which surrounds the filament (22). The authors also observed outer domain interactions in *Sinorhizobium meliloti* forming a dimeric outer domain interaction in two conformations (22). Similarly, *P. aeruginosa* PAO1 flagellin has been shown to form a ridged filament due to interactions of its outer domains (21). This suggests that along with the well-studied role of immune activation of the flagellin outer domains, there is also a role that these domains have on bacterial swim speed in different environments (25). In *S*. Typhimurium and *E. coli*, it has been previously shown that partial outer-domain-deleted FliC mutants are motile (45,46), however, full D2/3 deletions were non-motile, whereas our results show that *E. coli* K-12 expressing FliC deleted for domains D2 and D3 is still motile to some degree. This demonstrates that the outer domains of the *E. coli* K-12 flagellin are not absolutely required for motility. We observed rescue in motility when replacing the D2/3 domains with those from the *S*. Typhimurium flagellin as well as a two species chimera with *S*. Typhimurium and *Mesorhizobiium* sp. ORS 3359. Future structural work would characterise outer domain interactions in these mutant filaments quantify any effect on the superstructure of the filament.

The lack of motility in the majority of chimeric flagellin expressing strains could also arise from the lack of post-translational modifications. It is known that many flagellins, although not *E. coli* K-12, require glycosylation or methylation for the formation of the flagella filament (47,48). *Epsilonproteobacteria*, in which *H. mustelae* is found, require glycosylation for the formation of the flagellar filament (49). Both *P. aeruginosa* a-type and b-type flagellins also are glycosylated on their exposed surfaces (50). This may be the reason why no motility of some of the flagellin chimera expressing strains was observed. The levels of flagellin expression may also be an issue, compounded with complex flagellin structures that could potentially influence the formation of the flagellar filament. Previous work showed that motility was only partially restored (∼50%) when a rhamnose promoter was used when compared to the wild type flagellin promoter (51). Therefore, optimisation of the flagellin expression system, using various promoters and/or expression vectors, could increase the levels of chimeric flagellin expression and thus increase the likelihood of the formation of the flagellar filaments. Additionally, due to the use of multiple antibiotics to select and maintain the two expression plasmids there may have been a reduction in flagellin expression and therefore a reduced likelihood of functional filament formation (52).

Bacterial swim speed can vary widely with *E. coli* swim speeds ranging from 14.2-28.8 µm/s, *Bdellovibrio bacteriovorus* at 100 μm/s and *Thiovulum majus* at approximately 600 μm/s (53–56). *E. coli* and *T. majus* both contain flagellin outer domains whereas the *B. bacteriovorus* flagellin does not. We observed lower swim speeds and shorter filaments for EESS, EESM, EEE and EE expressing bacteria. Previous work has shown that even a single polymorphism (at position N87 in *E. coli* K-12 FliC) prevented motility in structured environments, although motility was observed in liquids (29). Therefore, it is plausible that whole domain replacements may alter the polymorphic transformation of the flagellar filament which allows for raised observable motility within soft agar. Additionally, reductions in motility may arise due to weaker filaments being more prone to shearing or due to differences in the filament export rate.

### Outer domain chimeras require *S*. Typhimurium D2 domain to form filaments

We observed that two out of eleven FliC variants rescued motility, with the remainder of the chimeras being non-motile. In work by Nedeljkocić *et al*., 13 flagellin chimeras, with combinations of all domains, were generated using the FliC from *S*. Typhimurium and *P. aeruginosa* PAO1 and only one multi-genus chimera produced a functional filament (21). Furthermore, in the same study the authors generated flagellin chimeras using domain swaps of two *P. aeruginosa* strains (PAO1 and PAK), where only two variants (where the D0 domain was exchanged) were motile (21). This further illustrates that the production of flagellin chimeras even within the same bacterial species does not yield a high proportion of motile variants. In *E. coli* SYC29 expressing chimeric-FliC containing outer domains, motility was only rescued only in chimeras that contained the *S*. Typhimurium D2 domain. This may indicate that there are key structural elements or residues found in this domain that allow of export and formation of a functional filament.

## Conclusion

Engineering of the flagellin outer domains can not only increase our understanding of filament formation but could also provide biotechnological benefits. Flagellin is already used to display antigens from other organisms and stimulate the immune system and has been engineered to contain fluorescent and enzymatic domains (25,57,58). Here we sought to develop chimeras for the outer domains of *E. coli* K-12 flagellin to provide evidence that recombination events have the potential to produce functional filaments. We have shown that two- and three-species chimeras can form functional filaments. However, our work illustrates the challenges of exchanging whole flagellin outer domains between bacteria. Further analysis is required to fully understand the export, folding and incorporation of chimeric flagellins and how they interact with other proteins within the bacterial flagellar motor. Future work will investigate how hypervariable FliC domains can affect motility in response to environmental changes and establish FliC and its homologs as a synthetic biology platform for biotechnological applications.

## Author Information

**Corresponding Author**

Correspondence to Matthew AB Baker

## References

1. Miyata M, Robinson RC, Uyeda TQ, Fukumori Y, Fukushima S ichi, Haruta S, et al. Tree of motility–A proposed history of motility systems in the tree of life. Genes to Cells. 2020;25(1):6– 21.

2. Sowa Y, Berry RM. Bacterial flagellar motor. Quarterly reviews of biophysics. 2008;41(2):103–32.

3. Nedeljković M, Sastre DE, Sundberg EJ. Bacterial Flagellar Filament: A Supramolecular Multifunctional Nanostructure. Int J Mol Sci. 2021 Jul 14;22(14).

4. Bulieris PV, Shaikh NH, Freddolino PL, Samatey FA. Structure of FlgK reveals the divergence of the bacterial Hook-Filament Junction of Campylobacter. Scientific reports. 2017;7(1):15743.

5. Hong HJ, Kim TH, Song WS, Ko HJ, Lee GS, Kang SG, et al. Crystal structure of FlgL and its implications for flagellar assembly. Scientific Reports. 2018;8(1):14307.

6. Postel S, Deredge D, Bonsor DA, Yu X, Diederichs K, Helmsing S, et al. Bacterial flagellar capping proteins adopt diverse oligomeric states. Elife. 2016;5:e18857.

7. Samatey FA, Imada K, Nagashima S, Vonderviszt F, Kumasaka T, Yamamoto M, et al. Structure of the bacterial flagellar protofilament and implications for a switch for supercoiling. Nature. 2001;410(6826):331–7.

8. Yonekura K, Maki-Yonekura S, Namba K. Complete atomic model of the bacterial flagellar filament by electron cryomicroscopy. Nature. 2003;424(6949):643–50.

9. Beatson SA, Minamino T, Pallen MJ. Variation in bacterial flagellins: from sequence to structure. Trends Microbiol. 2006 Apr;14(4):151–5.

10. O’Brien E, Bennett PM. Structure of straight flagella from a mutant Salmonella. Journal of molecular biology. 1972;70(1):133–52.

11. Kreutzberger MAB, Sonani RR, Liu J, Chatterjee S, Wang F, Sebastian AL, et al. Convergent evolution in the supercoiling of prokaryotic flagellar filaments. Cell. 2022 Sep 15;185(19):3487-3500.e14.

12. Végh BM, Gál P, Dobó J, Závodszky P, Vonderviszt F. Localization of the flagellum-specific secretion signal in Salmonella flagellin. Biochemical and biophysical research communications. 2006;345(1):93–8.

13. Yamaguchi T, Toma S, Terahara N, Miyata T, Ashihara M, Minamino T, et al. Structural and Functional Comparison of Salmonella Flagellar Filaments Composed of FljB and FliC. Biomolecules. 2020 Feb 6;10(2).

14. Il Kim M, Lee C, Park J, Jeon BY, Hong M. Crystal structure of Bacillus cereus flagellin and structure-guided fusion-protein designs. Sci Rep. 2018 Apr 11;8(1):5814.

15. Mohari B, Thompson MA, Trinidad JC, Setayeshgar S, Fuqua C. Multiple flagellin proteins have distinct and synergistic roles in Agrobacterium tumefaciens motility. Journal of Bacteriology. 2018;200(23):10–1128.

16. Wang F, Burrage AM, Postel S, Clark RE, Orlova A, Sundberg EJ, et al. A structural model of flagellar filament switching across multiple bacterial species. Nature communications. 2017;8(1):960.

17. Blum TB, Filippidou S, Fatton M, Junier P, Abrahams JP. The wild-type flagellar filament of the Firmicute Kurthia at 2.8 Å resolution in vivo. Scientific reports. 2019;9(1):14948.

18. San Martin F, Fule L, Iraola G, Buschiazzo A, Picardeau M. Diving into the complexity of the spirochetal endoflagellum. Trends in microbiology. 2023;31(3):294–307.

19. Hu D, Reeves PR. The Remarkable Dual-Level Diversity of Prokaryotic Flagellins. Zhaxybayeva O, editor. mSystems. 2020 Feb 11;5(1):e00705–19.

20. Fields JL, Zhang H, Bellis NF, Petersen HA, Halder SK, Rich-New ST, et al. Structural diversity and clustering of bacterial flagellar outer domains. Nat Commun. 2024 Nov 3;15:9500.

21. Nedeljković M, Kreutzberger MA, Postel S, Bonsor D, Xing Y, Jacob N, et al. An unbroken network of interactions connecting flagellin domains is required for motility in viscous environments. PLoS Pathogens. 2023;19(5):e1010979.

22. Kreutzberger MA, Sobe RC, Sauder AB, Chatterjee S, Peña A, Wang F, et al. Flagellin outer domain dimerization modulates motility in pathogenic and soil bacteria from viscous environments. Nature communications. 2022;13(1):1422.

23. Esteves NC, Bigham DN, Scharf BE. Phages on filaments: A genetic screen elucidates the complex interactions between Salmonella enterica flagellin and bacteriophage Chi. PLoS Pathogens. 2023;19(8):e1011537.

24. Eaves-Pyles TD, Wong HR, Odoms K, Pyles RB. Salmonella flagellin-dependent proinflammatory responses are localized to the conserved amino and carboxyl regions of the protein. The Journal of Immunology. 2001;167(12):7009–16.

25. Hajam IA, Dar PA, Shahnawaz I, Jaume JC, Lee JH. Bacterial flagellin—a potent immunomodulatory agent. Experimental & molecular medicine. 2017;49(9):e373–e373.

26. Liu G, Tarbet B, Song L, Reiserova L, Weaver B, Chen Y, et al. Immunogenicity and efficacy of flagellin-fused vaccine candidates targeting 2009 pandemic H1N1 influenza in mice. PloS one. 2011;6(6):e20928.

27. Taylor DN, Treanor JJ, Strout C, Johnson C, Fitzgerald T, Kavita U, et al. Induction of a potent immune response in the elderly using the TLR-5 agonist, flagellin, with a recombinant hemagglutinin influenza–flagellin fusion vaccine (VAX125, STF2. HA1 SI). Vaccine. 2011;29(31):4897–902.

28. Datsenko KA, Wanner BL. One-step inactivation of chromosomal genes in Escherichia coli K-12 using PCR products. Proceedings of the National Academy of Sciences. 2000 Jun 6;97(12):6640– 5.

29. Kinosita Y, Ishida T, Yoshida M, Ito R, Morimoto YV, Goto K, et al. Distinct chemotactic behavior in the original Escherichia coli K-12 depending on forward-and-backward swimming, not on run-tumble movements. Sci Rep. 2020 Sep 28;10(1):15887.

30. UniProt: the universal protein knowledgebase in 2023. Nucleic acids research. 2023;51(D1):D523–31.

31. Suzek BE, Wang Y, Huang H, McGarvey PB, Wu CH, UniProt Consortium. UniRef clusters: a comprehensive and scalable alternative for improving sequence similarity searches. Bioinformatics. 2015;31(6):926–32.

32. Miller MA, Pfeiffer W, Schwartz T. Creating the CIPRES Science Gateway for inference of large phylogenetic trees. In Ieee; 2010. p. 1–8.

33. Edgar RC. MUSCLE: multiple sequence alignment with high accuracy and high throughput. Nucleic acids research. 2004;32(5):1792–7.

34. Larsson A. AliView: a fast and lightweight alignment viewer and editor for large datasets. Bioinformatics. 2014;30(22):3276–8.

35. Nguyen LT, Schmidt HA, Von Haeseler A, Minh BQ. IQ-TREE: a fast and effective stochastic algorithm for estimating maximum-likelihood phylogenies. Molecular biology and evolution. 2015;32(1):268–74.

36. Price MN, Dehal PS, Arkin AP. FastTree 2 – Approximately Maximum-Likelihood Trees for Large Alignments. Poon AFY, editor. PLoS ONE. 2010 Mar 10;5(3):e9490.

37. Letunic I, Bork P. Interactive Tree of Life (iTOL) v6: recent updates to the phylogenetic tree display and annotation tool. Nucleic Acids Research. 2024 Jul 5;52(W1):W78–82.

38. Abramson J, Adler J, Dunger J, Evans R, Green T, Pritzel A, et al. Accurate structure prediction of biomolecular interactions with AlphaFold 3. Nature. 2024 Jun;630(8016):493–500.

39. Ishida T, Ito R, Clark J, Matzke NJ, Sowa Y, Baker MA. Sodium-powered stators of the bacterial flagellar motor can generate torque in the presence of phenamil with mutations near the peptidoglycan-binding region. Molecular Microbiology. 2019;111(6):1689–99.

40. Schindelin J, Arganda-Carreras I, Frise E, Kaynig V, Longair M, Pietzsch T, et al. Fiji: an open-source platform for biological-image analysis. Nature methods. 2012;9(7):676–82.

41. Ridone P, Winter DL, Baker MAB. Tuning the stator subunit of the flagellar motor with coiled-coil engineering. Protein Science. 2023;32(12):e4811.

42. Kreutzberger MAB, Ewing C, Poly F, Wang F, Egelman EH. Atomic structure of the Campylobacter jejuni flagellar filament reveals how ε Proteobacteria escaped Toll-like receptor 5 surveillance. Proc Natl Acad Sci U S A. 2020 Jul 21;117(29):16985–91.

43. Poggio S, Abreu-Goodger C, Fabela S, Osorio A, Dreyfus G, Vinuesa P, et al. A Complete Set of Flagellar Genes Acquired by Horizontal Transfer Coexists with the Endogenous Flagellar System in Rhodobacter sphaeroides. J Bacteriol. 2007 Apr;189(8):3208–16.

44. Linton D, Hurtado A, Lawson AJ, Clewley JP, Chart H, Stanley J. Campylobacter coli strains with enlarged flagellin genes isolated from river water. Res Microbiol. 1999 May;150(4):247–55.

45. Malapaka RRV, Adebayo LO, Tripp BC. A deletion variant study of the functional role of the Salmonella flagellin hypervariable domain region in motility. Journal of molecular biology. 2007;365(4):1102–16.

46. Kuwajima G. Construction of a minimum-size functional flagellin of Escherichia coli. J Bacteriol. 1988 Jul;170(7):3305–9.

47. De Maayer P, Cowan DA. Comparative genomic analysis of the flagellin glycosylation island of the Gram-positive thermophile Geobacillus. BMC Genomics. 2016 Nov 14;17(1):913.

48. Horstmann JA, Lunelli M, Cazzola H, Heidemann J, Kühne C, Steffen P, et al. Methylation of Salmonella Typhimurium flagella promotes bacterial adhesion and host cell invasion. Nat Commun. 2020 Apr 24;11:2013.

49. Thibault P, Logan SM, Kelly JF, Brisson JR, Ewing CP, Trust TJ, et al. Identification of the carbohydrate moieties and glycosylation motifs in Campylobacter jejuni flagellin. J Biol Chem. 2001 Sep 14;276(37):34862–70.

50. Verma A, Schirm M, Arora SK, Thibault P, Logan SM, Ramphal R. Glycosylation of b-Type Flagellin of Pseudomonas aeruginosa: Structural and Genetic Basis. J Bacteriol. 2006 Jun;188(12):4395– 403.

51. Thomson NM, Pallen MJ. Restoration of wild-type motility to flagellin-knockout Escherichia coli by varying promoter, copy number and induction strength in plasmid-based expression of flagellin. Curr Res Biotechnol. 2020 Nov;2:45–52.

52. San Millan A, MacLean RC. Fitness Costs of Plasmids: a Limit to Plasmid Transmission. Microbiol Spectr. 5(5): 10.1128/microbiolspec.mtbp-0016–2017.

53. Berg HC, Brown DA. Chemotaxis in Escherichia coli analysed by three-dimensional tracking. nature. 1972;239(5374):500–4.

54. Macnab RM, Koshland DE. The Gradient-Sensing Mechanism in Bacterial Chemotaxis. Proc Natl Acad Sci U S A. 1972 Sep;69(9):2509–12.

55. Herzog B, Wirth R. Swimming Behavior of Selected Species of Archaea. Appl Environ Microbiol. 2012 Mar 15;78(6):1670–4.

56. Fenchel T. Motility and chemosensory behaviour of the sulphur bacterium Thiovulum majus. Microbiology. 1994 Nov 1;140(11):3109–16.

57. Klein A, Tóth B, Jankovics H, Muskotál A, Vonderviszt F. A polymerizable GFP variant. Protein Engineering, Design & Selection. 2012;25(4):153–7.

58. Klein Á, Szabó V, Kovács M, Patkó D, Tóth B, Vonderviszt F. Xylan-Degrading Catalytic Flagellar Nanorods. Mol Biotechnol. 2015 Sep;57(9):814–9.

