## Supplementary material for "Rescue of bacterial motility using two and three-species FliC chimeras": Supplmentary Tables S1-S5, Supplementary Figures S1-S5.

### Tables S1-5

### Figures S1-S5

Jacob Scadden[1], Pietro Ridone[1], Divyangi Pandit[1], Yoshiyuki Sowa[2,3], Matthew AB Baker[1*]

[1]School of Biotechnology and Biomolecular Sciences, University of New South Wales, Sydney, Australia
[2]Department of Frontier Bioscience, Hosei University, Tokyo, Japan
[3]Research Center for Micro-Nano Technology, Hosei University, Tokyo, Japan

*Corresponding Author

**Table S1**. Bacterial strains and plasmids used in this study.

| **Strain** | **Relevant Characteristics** | **Reference** |
|---|---|---|
| *E. coli* NEB 10-β | DH10B derivative. Shuttle vector cloning strain | New England Biolabs (C3019) |
| *E. coli* SYC29 | *E. coli* RP437, Δ*motAB fliC*::*tetRA*, TET$^{R}$ | This study |
| **Plasmid** | **Relevant Characteristics** | **Reference** |
| pET21(+) | Expression vector, T7 promoter, Amp$^{R}$, IPTG induction | Merck (69740) |
| pDB108 | pBAD33 backbone, motAB, CAM$^{R}$, arabinose induction | David Blair* (University of Utah, USA) |
| pET21_*EEEE* | Wild type E. coli *fliC*, pET21a backbone | This study |
| pET21_*EEE* | *E. coli fliC* with deletion of 195-300, pET21a backbone | This study |
| pET21_*EE* | *fliC* with deletion of 174-405, pET21a backbone | This study |
| pET21_*EESS* | *E. coli fliC* with D2/3 domains replaced by *Salmonella* Typhimurium strain LT2 D2/3 domains, pET21a backbone | This study |
| pET21_*EESM* | *E. coli fliC* with D2/3 domains replaced by *Salmonella* Typhimurium strain LT2 D2 domain and the *Mesorhizobium* sp. ORS 3359 D2 domain, pET21a backbone | This study |
| pET21_*EESP* | *E. coli fliC* with D2/3 domains replaced by *Salmonella* Typhimurium strain LT2 D2 domain and the *Pseudomonas aeruginosa* D3 domain, pET21a backbone | This study |
| pET21_*EESC* | *E. coli fliC* with D2/3 domains replaced by *Salmonella* | This study |

| | | |
|---|---|---|
| | Typhimurium strain LT2 D2 domain and the *Collimonas fungivorans* strain Ter331 D2 domain, pET21a backbone | |
| pET21_*EEHH* | *E. coli fliC* with D2/3 domains replaced by the *Helicobacter mustelae* D2/3 domains, pET21a backbone | This study |
| pET21_*EEC* | *E. coli fliC* with D2/3 domains replaced by the *Collimonas fungivorans* strain Ter331 D2 domains, pET21a backbone | This study |
| pET21_EEES | *E. coli fliC* with the D3 domain replaced by the *Salmonella* Typhimurium strain LT2 D3 domain, pET21a backbone | This Study |
| pET21_EEEM | *E. coli fliC* with the D3 domain replaced by the *Mesorhizobium* sp. ORS 3359 D2 domain, pET21a backbone | This Study |
| pET21_EEEC | *E. coli fliC* with the D3 domain replaced by the *Collimonas fungivorans* strain Ter331 D2 domain, pET21a backbone | This Study |
| pET21_EEEP | *E. coli fliC* with the D3 domain replaced by the *Pseudomonas aeruginosa* D3 domain, pET21a backbone | This Study |
| pET21_EEEH | *E. coli fliC* with the D3 domain replaced by the *Helicobacter mustelae* D3 domain, pET21a backbone | This Study |

Kindly gifted from the *Blair Lab

**Table S2**. Flagellin reference sequences used in the BLASTP search. The final row, FliD, (**Bold**) was used as the outgroup (P24216).

| **UniProt ID** | **Bacteria** | **Protein ID** | **Length (Amino Acids)** | **D2/3 Domains Present** |
|---|---|---|---|---|
| P04949 | *E. coli* (strain K12) | Flagellin | 498 | Yes |
| A0A1H7N2K9 | *Blastococcus* sp. DSM 46786 | Flagellin | 277 | No |
| P13118 | *Rhizobium meliloti* | Flagellin A | 395 | D2 Only |
| Q06064 | *Bordetella bronchiseptica* (strain ATCC BAA-588) | Flagellin | 391 | D2 Only |
| Q6AJQ8 | *Desulfotalea psychrophile* (DSM 12343) | Flagellin | 857 | Yes |
| P0A0S1 | *Helicobacter pylori* (strain ATCC 700392) | Flagellin A | 510 | Yes |
| P80583 | *Clostridium tyrobutyricum* | Flagellin | 377 | D2 Only |

| | | | | |
|---|---|---|---|---|
| A0A1N6WG11 | *Alkalispirochaeta americana* | Flagellin | 284 | No |
| **P24216** | ***E. coli* (strain K12)** | **Flagellar hook-associated protein** | **468** | **N/A** |

**Table S3**. Parameters used for the generation of the flagellin phylogeny using IQTree on ACCESS.

| IQ-Tree on ACCESS - Parameters | |
|---|---|
| abayes_test_ | FALSE |
| datatype_ | PROTEIN |
| extract_datefile_ | FALSE |
| fix_branchlengths_ | FALSE |
| lbp_test_ | FALSE |
| median_approximation_ | FALSE |
| more_memory2_ | FALSE |
| more_memory_ | FALSE |
| no_mlpairwise_ | FALSE |
| optimize_weights_ | FALSE |
| parametrical_test_ | FALSE |
| per_sitefile_ | FALSE |
| per_sitewplfile_ | FALSE |
| per_sitewslfile_ | FALSE |
| per_sitewslmfile_ | FALSE |
| per_sitewslmrfile_ | FALSE |
| per_sitewslrfile_ | FALSE |
| per_sitewspmfile_ | FALSE |
| per_sitewspmrfile_ | FALSE |
| per_sitewsprfile_ | FALSE |
| print_sitestats_ | FALSE |
| runtime_ | 0.5 |
| sh_test_ | FALSE |
| slower_NNI_ | FALSE |
| specify_fulltreesearch_ | FALSE |
| specify_numpatterns_ | 921 |
| specify_prefix_ | output |
| specify_radius_ | 6 |
| specify_safe_ | FALSE |
| specify_symtype_ | pval |
| thorough_estimation_ | FALSE |
| unbiased_test_ | FALSE |
| use_fasttreesearch_ | FALSE |
| use_nj_ | FALSE |
| use_random_ | FALSE |
| use_symmtest_ | FALSE |

| | |
|---|---|
| which_iqtree_ | 232 |
| write_ancestralseqs_ | FALSE |
| write_locally_optimal_ | FALSE |
| write_loglikelihoods_ | FALSE |
| write_sitelikelihoods_ | FALSE |

**Table S4**. Sequences of FliC chimeras and outer domain deleted FliC. Sequences are coloured based on domains incorporated into the outer domains for *E. coli* K-12 FliC.

| FliC Variant | Domains Present | Amino Acid Sequence | Length |
|---|---|---|---|
| EEEE | D0, D1, D2, D3 | MAQVINTNSLSLITQNNINKNQSALSSSIERLSSGLRINSAKDDAAGQAIANRFTSNIKGLTQAARNANDGISVAQTTEGALSEINNNLQRVRELTVQATTGTNSESDLSSIQDEIKSRLDEIDRVSGQTQFNGVNVLAKNGSMKIQVGANDNQTITIDLKQIDAKTLGLDGFSVKNNDTVTTSAPVTAFGATTTNNIKLTGITLSTEAATDTGGTNPASIEGVYTDNGNDYYAKITGGDNDGKYYAVTVANDGTVTMATGATANATVTDANTTKATTITSGGTPVQIDNTAGSATANLGAVSLVKLQDSKGNDTDTYALKDTNGNLYAADVNETTGAVSVKTITYTDSSGAASSPTAVKLGGDDGKTEVVDIDGKTYDSADLNGGNLQTGLTAGGEALTAVANGKTTDPLKALDDAIASVDKFRSSLGAVQNRLDSAVTNLNNTTTNLSEAQSRIQDADYATEVSNMSKAQIIQQAGNSVLAKANQVPQQVLSLLQG | 498 |
| EEE | D0, D1, D2 | MAQVINTNSLSLITQNNINKNQSALSSSIERLSSGLRINSAKDDAAGQAIANRFTSNIKGLTQAARNANDGISVAQTTEGALSEINNNLQRVRELTVQATTGTNSESDLSSIQDEIKSRLDEIDRVSGQTQFNGVNVLAKNGSMKIQVGANDNQTITIDLKQIDAKTLGLDGFSVKNNDTVTTSAPVTAFGATTAVSLVKLQDSKGNDTDTYALKDTNGNLYAADVNETTGAVSVKTITYTDSSGAASSPTAVKLGGDDGKTEVVDIDGKTYDSADLNGGNLQTGLTAGGEALTAVANGKTTDPLKALDDAIASVDKFRSSLGAVQNRLDSAVTNLNNTTTNLSEAQSRIQDADYATEVSNMSKAQIIQQAGNSVLAKANQVPQQVLSLLQG | 393 |
| EE | D0, D1 | MAQVINTNSLSLITQNNINKNQSALSSSIERLSSGLRINSAKDDAAGQAIANRFTSNIKGLTQAARNANDGISVAQTTEGALSEINNNLQRVRELTVQATTGTNSESDLSSIQDEIKSRLDEIDRVSGQTQFNGVNVLAKNGSMKIQVGANDNQTITIDLKQIDAKTLGLDGFKTTDPLKALDDAIASVDKFRSSLGAVQNRLDSAVTNLNNTTTNLSEAQSRIQDADYATEVSNMSKAQIIQQAGNSVLAKANQVPQQVLSLLQG | 266 |
| EEHH | D0, D1, D2, D3 | MAQVINTNSLSLITQNNINKNQSALSSSIERLSSGLRINSAKDDAAGQAIANRFTSNIKGLTQAARNANDGISVAQTTEGALSEINNNLQRVRELTVQATTGTNSESDLSSIQDEIKSRLDEIDRVSGQTQFNGVNVLAKNGSMKIQVGANDNQTITIDLKQIDAKTLGLDGFQVRINTGAMITAASEATLTFKQINGGGTSPLEGVKISHSVGTGLGVLAEVINKNSDKTGIRAKASVETTSDKEIMSGNLKNLTINDVNIGNIVDIKKGDADGRLVQAINALTSSTGVEASTDSKGRLNLRSVDGRGIVLKADASEDNGDGKSAPMAIDAVNGGQSITDGEGAANYGRLSLVRLDARDIVLTSSDKPDENKFSAIGFGDNNVAMATVNLRDVLGKFDASVKSASGANYNAVIASGNSNLKTTDPLKALDDAIASVDKFRSSLGAVQNRLDSAVTNLNNTTTNLSEAQSRIQDADYATEVSNMSKAQIIQQAGNSVLAKANQVPQQVLSLLQG | 514 |
| EEC | D0, D1, D2 | MAQVINTNSLSLITQNNINKNQSALSSSIERLSSGLRINSAKDDAAGQAIANRFTSNIKGLTQAARNANDGISVAQTTEGALSEINNNLQRVRELTVQATTGTNSESDLSSIQDEIKSRLDEIDRVSGQTQFNGVNVLAKNGSMKIQVGANDNQTITIDLKQIDAKTLGLDGFNTLNLAGPSGATAAMTPTGKDNTGASVAVAAGGFTAVNRDAAAGFTATVADLVTDARGNTYVKNGANFYAATVTVADATQGATAATPTATVTYDSSKSLTAASGSPLKTTDPLKALDDAIASVDKFRSSLGAVQNRLDSAVTNLNNTTTNLSEAQSRIQDADYATEVSNMSKAQIIQQAGNSVLAKANQVPQQVLSLLQG | 373 |
| EESS | D0, D1, D2, D3 | MAQVINTNSLSLITQNNINKNQSALSSSIERLSSGLRINSAKDDAAGQAIANRFTSNIKGLTQAARNANDGISVAQTTEGALSEINNNLQRVRELTVQATTGTNSESDLSSIQDEIKSRLDEIDRVSGQTQFNGVNVLAKNGSMKIQVGANDNQTITIDLKQIDAKTLGLDGFQKYKVSDTAATVTG | 499 |

| | | | |
|---|---|---|---|
| | | YADTTIALDNSTFKASATGLGGTDQKIDGDLKFDDTTGKYYAKVTVTGGTGKDGYYEVSVDKTNGEVTLAGGATSPLTGGLPATATEDVKNVQVANADLTEAKAALTAAGVTGTASVVKMSYTDNNGKTIDGGLAVKVGDDYYSATQNKDGSISINTTKYTADDGTSKTALNKLGGADGKTEVVSIGGKTYAASKAEGHNFKAQPDLAEAAATTTENPLKTTDPLKALDDAIASVDKFRSSLGAVQNRLDSAVTNLNNTTTNLSEAQSRIQDADYATEVSNMSKAQIIQQAGNSVLAKANQVPQQVLSLLQG | |
| EESM | D0, D1, D2, D2 | MAQVINTNSLSLITQNNINKNQSALSSSIERLSSGLRINSAKDDAAGQAIANRFTSNIKGLTQAARNANDGISVAQTTEGALSEINNNLQRVRELTVQATTGTNSESDLSSIQDEIKSRLDEIDRVSGQTQFNGVNVLAKNGSMKIQVGANDNQTITIDLKQIDAKTLGLDGFQKYKVSDTAATVTINVESTKLFDSGLSTAVKNQGILDRKTSIYSTAADQSAYDAAYATAIAGGATDIAANTAGQAAVAGTPVKIDNVSAFNLDITATGVTDDIITQMVTNADLTEAKAALTAAGVTGTASVVKMSYTDNNGKTIDGGLAVKVGDDYYSATQNKDGSISINTTKYTADDGTSKTALNKLGGADGKTEVVSIGGKTYAASKAEGHNFKAQPDLAEAAATTTENPLKTTDPLKALDDAIASVDKFRSSLGAVQNRLDSAVTNLNNTTTNLSEAQSRIQDADYATEVSNMSKAQIIQQAGNSVLAKANQVPQQVLSLLQG | 499 |
| EESC | D0, D1, D2, D2 | MAQVINTNSLSLITQNNINKNQSALSSSIERLSSGLRINSAKDDAAGQAIANRFTSNIKGLTQAARNANDGISVAQTTEGALSEINNNLQRVRELTVQATTGTNSESDLSSIQDEIKSRLDEIDRVSGQTQFNGVNVLAKNGSMKIQVGANDNQTITIDLKQIDAKTLGLDGFQKYKVSDTAATVTNTLNLAGPSGATAAMTPTGKDNTGASVAVAAGGFTAVNRDAAAGFTATVADLVTDARGNTYVKNGANFYAATVTVADATQGATAATPTATVTYDSSKSLTAASGSPLNADLTEAKAALTAAGVTGTASVVKMSYTDNNGKTIDGGLAVKVGDDYYSATQNKDGSISINTTKYTADDGTSKTALNKLGGADGKTEVVSIGGKTYAASKAEGHNFKAQPDLAEAAATTTENPLKTTDPLKALDDAIASVDKFRSSLGAVQNRLDSAVTNLNNTTTNLSEAQSRIQDADYATEVSNMSKAQIIQQAGNSVLAKANQVPQQVLSLLQG | 510 |
| EESP | D0, D1, D2, D2 | MAQVINTNSLSLITQNNINKNQSALSSSIERLSSGLRINSAKDDAAGQAIANRFTSNIKGLTQAARNANDGISVAQTTEGALSEINNNLQRVRELTVQATTGTNSESDLSSIQDEIKSRLDEIDRVSGQTQFNGVNVLAKNGSMKIQVGANDNQTITIDLKQIDAKTLGLDGFQKYKVSDTAATVTNGTYFKADGGGAVTAATASGTVDIAIGITGGSAVNVKVDMKGNETAEQAAAKIAAAVNDANVGIGAFSDGDTISYVSKAGKDGSGAITSAVSGVVIADTGSTGVGTAAGVTPSATAFAKTNDTVAKIDISTANADLTEAKAALTAAGVTGTASVVKMSYTDNNGKTIDGGLAVKVGDDYYSATQNKDGSISINTTKYTADDGTSKTALNKLGGADGKTEVVSIGGKTYAASKAEGHNFKAQPDLAEAAATTTENPLKTTDPLKALDDAIASVDKFRSSLGAVQNRLDSAVTNLNNTTTNLSEAQSRIQDADYATEVSNMSKAQIIQQAGNSVLAKANQVPQQVLSLLQG | 535 |
| EEES | D0, D1, D2, D3 | MAQVINTNSLSLITQNNINKNQSALSSSIERLSSGLRINSAKDDAAGQAIANRFTSNIKGLTQAARNANDGISVAQTTEGALSEINNNLQRVRELTVQATTGTNSESDLSSIQDEIKSRLDEIDRVSGQTQFNGVNVLAKNGSMKIQVGANDNQTITIDLKQIDAKTLGLDGFSVKNNDTVTTSAPVTGYADTTIALDNSTFKASATGLGGTDQKIDGDLKFDDTTGKYYAKVTVTGGTGKDGYYEVSVDKTNGEVTLAGGATSPLTGGLPATATEDVKNVQVAAFGATTAVSLVKLQDSKGNDTDTYALKDTNGNLYAADVNETTGAVSVKTITYTDSSGAASSPTAVKLGGDDGKTEVVDIDGKTYDSADLNGGNLQTGLTAGGEALTAVANGKTTDPLKALDDAIASVDKFRSSLGAVQNRLDSAVTNLNNTTTNLSEAQSRIQDADYATEVSNMSKAQIIQQAGNSVLAKANQVPQQVLSLLQG | 488 |
| EEEM | D0, D1, D2, D2 | MAQVINTNSLSLITQNNINKNQSALSSSIERLSSGLRINSAKDDAAGQAIANRFTSNIKGLTQAARNANDGISVAQTTEGALSEINNNLQRVRELTVQATTGTNSESDLSSIQDEIKSRLDEIDRVSGQTQFNGVNVLAKNGSMKIQVGANDNQTITIDLKQIDAKTLGLDGFSVKNNDTVTTSAPVTINVESTKLFDSGLSTAVKNQGILDRKTSIYSTAADQSAYDAAYATAIAGGATDIAANTAGQAAVAGTPVKIDNVSAFNLDITATGVTDDIITQMVTAFGATTAVSLVKLQDSKGNDTDTYALKDTNGNLYAADVNETTGAVSVKTITYTDSSGAASSPTAVKLGGDDGKTEVVDIDGKTYDSADLNGGNLQTGLTAGGEALTAVANGKTTDPLKALDDAIASVDKFRSSLGAVQNRLDSAVTNLNNTTTNLSEAQSRIQDADYATEVSNMSKAQIIQQAGNSVLAKANQVPQQVLSLLQG | 488 |
| EEEC | D0, D1, D2, D2 | MAQVINTNSLSLITQNNINKNQSALSSSIERLSSGLRINSAKDDAAGQAIANRFTSNIKGLTQAARNANDGISVAQTTEGALSEINNNLQRVRELTVQATTGTNSESDLSSIQDEIKSRLDEIDRVSGQTQFNGVNVLAKNGSMKIQVGANDNQTITIDLKQIDAKTLGLDGFSVKNNDTVTTSAPV | 489 |

| | | | |
|---|---|---|---|
| | | TNTLNLAGPSGATAAMTPTGKDNTGASVAVAAGGFTAVNRDAAAGFTATVADLVTDARGNTYVKNGANFYAATVTVADATQGATAATPTATVTYDSSKSLTAASGSPLAFGATTAVSLVKLQDSKGNDTDTYALKDTNGNLYAADVNETTGAVSVKTITYTDSSGAASSPTAVKLGGDDGKTEVVDIDGKTYDSADLNGGNLQTGLTAGGEALTAVANGKTTDPLKALDDAIASVDKFRSSLGAVQNRLDSAVTNLNNTTTNLSEAQSRIQDADYATEVSNMSKAQIIQQAGNSVLAKANQVPQQVLSLLQG | |
| EEEP | D0, D1, D2, D3 | MAQVINTNSLSLITQNNINKNQSALSSSIERLSSGLRINSAKDDAAGQAIANRFTSNIKGLTQAARNANDGISVAQTTEGALSEINNNLQRVRELTVQATTGTNSESDLSSIQDEIKSRLDEIDRVSGQTQFNGVNVLAKNGSMKIQVGANDNQTITIDLKQIDAKTLGLDGFSVKNNDTVTTSAPVTNGTYFKADGGGAVTAATASGTVDIAIGITGGSAVNVKVDMKGNETAEQAAAKIAAAVNDANVGIGAFSDGDTISYVSKAGKDGSGAITSAVSGVVIADTGSTGVGTAAGVTPSATAFAKTNDTVAKIDISTAAFGATTAVSLVKLQDSKGNDTDTYALKDTNGNLYAADVNETTGAVSVKTITYTDSSGAASSPTAVKLGGDDGKTEVVDIDGKTYDSADLNGGNLQTGLTAGGEALTAVANGKTTDPLKALDDAIASVDKFRSSLGAVQNRLDSAVTNLNNTTTNLSEAQSRIQDADYATEVSNMSKAQIIQQAGNSVLAKANQVPQQVLSLLQG | 522 |
| EEEH | D0, D1, D2, D3 | MAQVINTNSLSLITQNNINKNQSALSSSIERLSSGLRINSAKDDAAGQAIANRFTSNIKGLTQAARNANDGISVAQTTEGALSEINNNLQRVRELTVQATTGTNSESDLSSIQDEIKSRLDEIDRVSGQTQFNGVNVLAKNGSMKIQVGANDNQTITIDLKQIDAKTLGLDGFSVKNNDTVTTSAPVTDKEIMSGNLKNLTINDVNIGNIVDIKKGDADGRLVQAINALTSSTGVEASTDSKGRLNLRSVDGRGIVLKADASEDNGDGKSAPMAIDAVNGGQSITDGEGAAFGATTAVSLVKLQDSKGNDTDTYALKDTNGNLYAADVNETTGAVSVKTITYTDSSGAASSPTAVKLGGDDGKTEVVDIDGKTYDSADLNGGNLQTGLTAGGEALTAVANGKTTDPLKALDDAIASVDKFRSSLGAVQNRLDSAVTNLNNTTTNLSEAQSRIQDADYATEVSNMSKAQIIQQAGNSVLAKANQVPQQVLSLLQG | 492 |

**Table S5**. Primers and PCR conditions used in this study to confirm chimeric and outer domain deleted *fliC* into pET21(+).

| Primer Name | Template Sequence | 5' -> 3' Sequence | Tm (°C) | Length |
|---|---|---|---|---|
| pET21_F | pET21(+) | TTCCTTTCGGGCTTTGTTAGCAG | 66.4 | 23 |
| pET21_R | pET21(+) | GGATCGAGATCTCGATCCCGC | 66.9 | 21 |

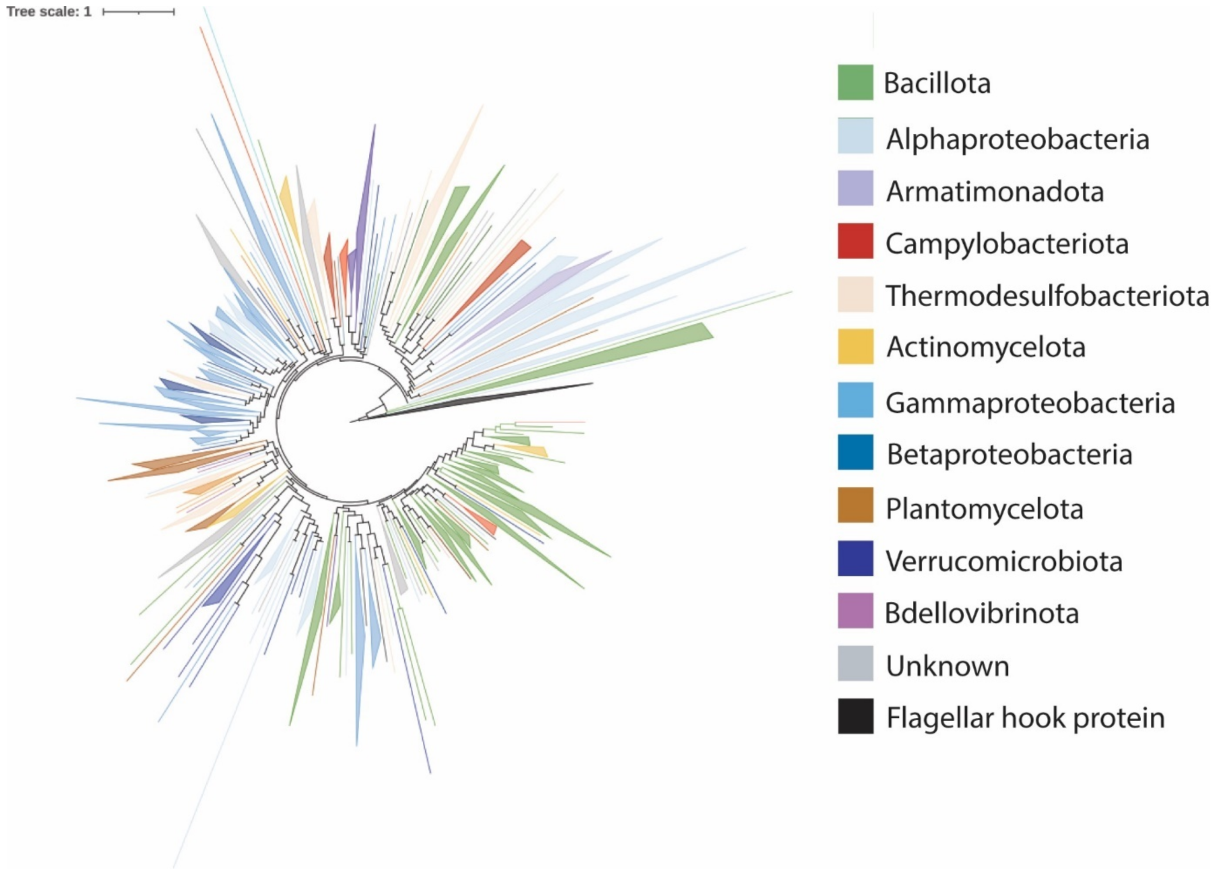

**Figure S1**. Phylogenetic tree generated using FastTree of 605 available representative flagellin sequences from Fields *et al*., (20). 14 hook-associated protein sequences as the root. Clades were determined if two or more branches from the same phylum/family were adjacent.

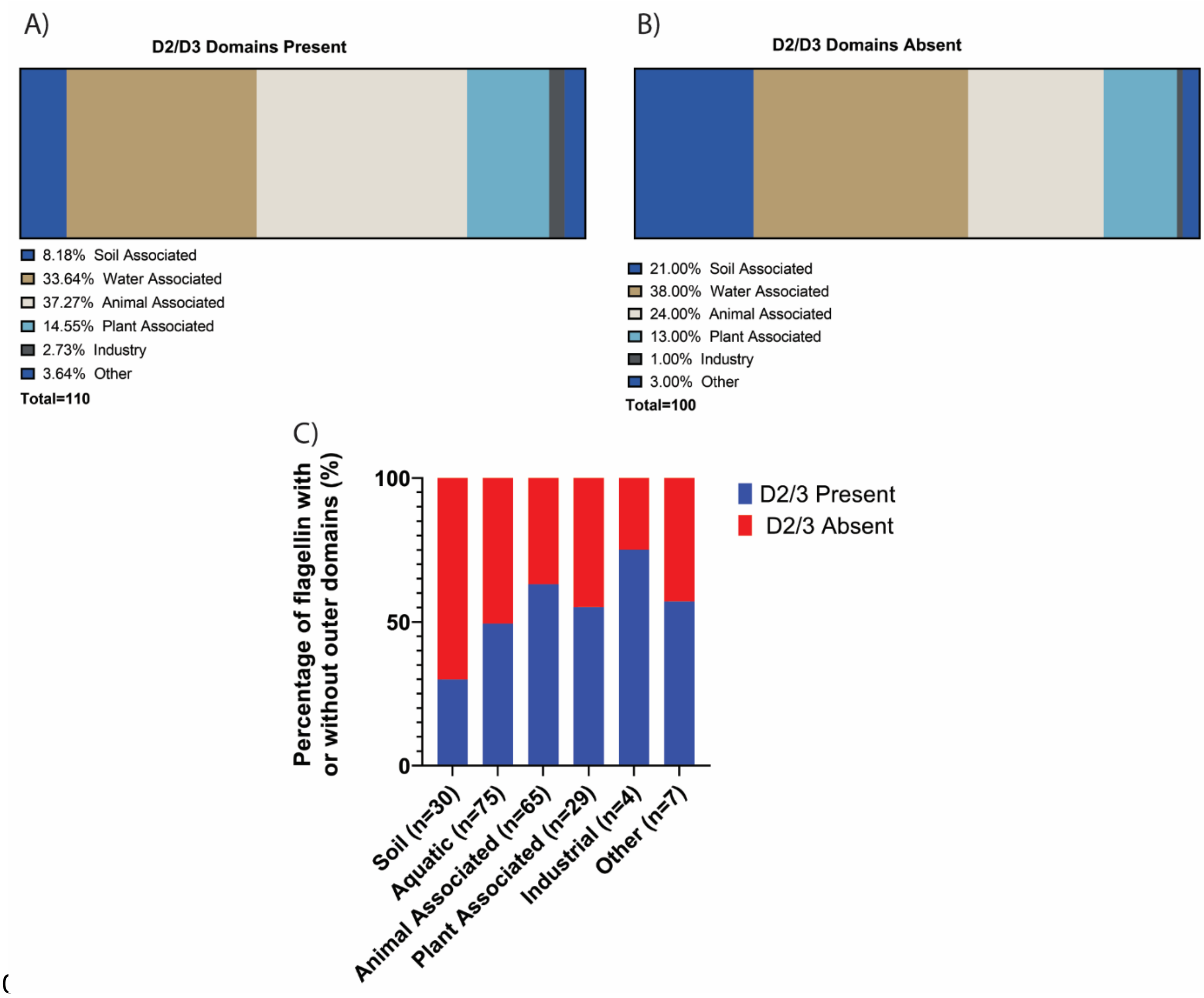

**Figure S2**. Isolation sources of bacterial flagellins with outer domains A) present and B) absent used to make the flagellin phylogeny. Isolation sources are group in six major clusters; Soil (Blue), Aquatic (Dark Brown), Animal Associated (Light Brown), Plant Associated (Light Blue), Industrial (Grey) and Other (Dark Blue). C) Percentages of flagellins with and without outer domains found from each isolation source.

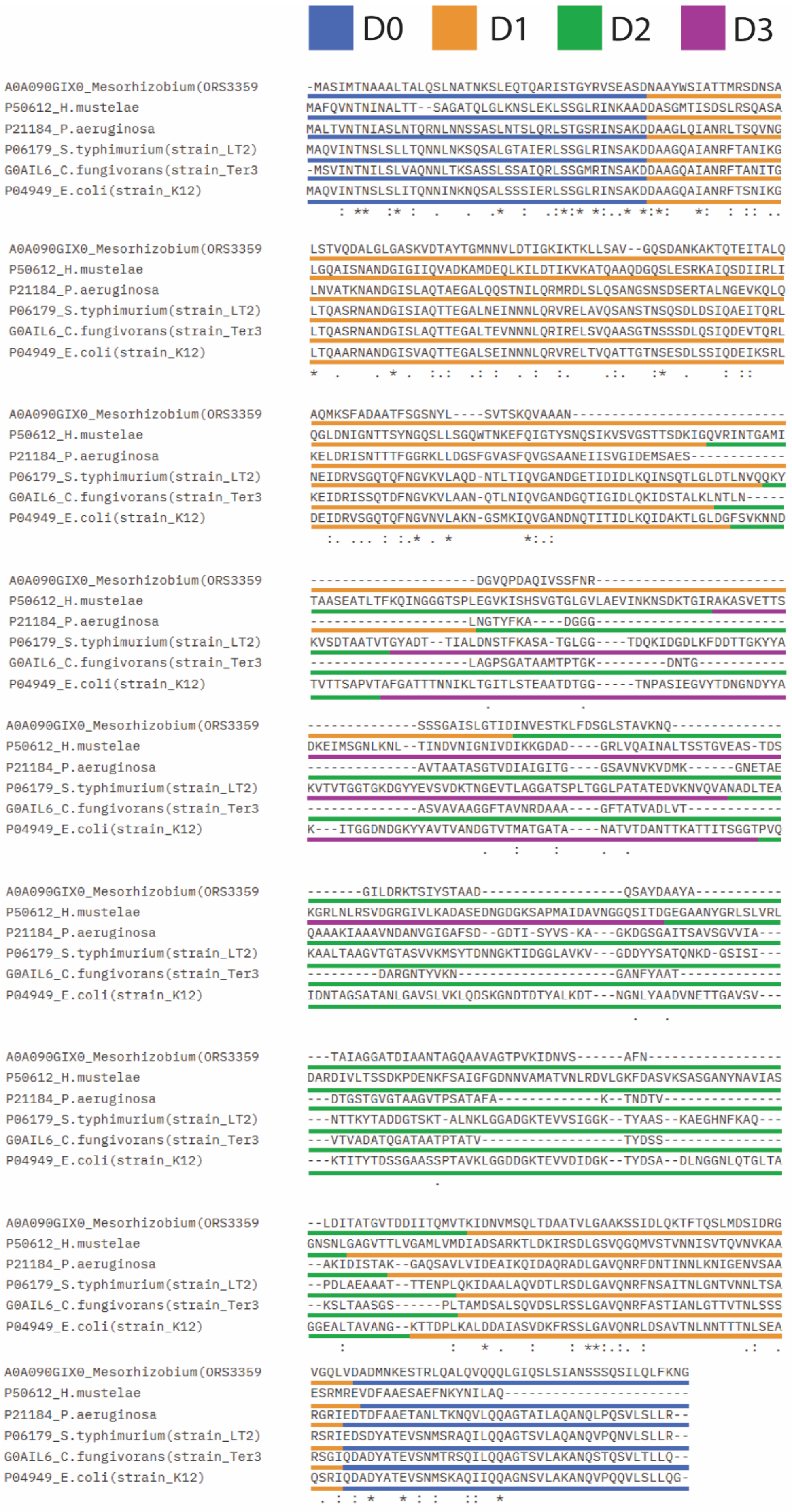
D0
D1
D2
D3
A0A090GIX0_Mesorhizobium(ORS3359 -MASIMTNAAALTALQSLNATNKSLEQTQARISTGYRVSEASDNAAYWSIATTMRSDNSA
P50612_H.mustelae MAFQVNTNINALTT--SAGATQLGLKNSLEKLSSGLRINKAADDASGMTISDSLRSQASA
P21184_P.aeruginosa MALTVNTNIASLNTQRNLNNSSASLNTSLQRLSTGSRINSAKDDAAGLQIANRLTSQVNG
P06179_S.typhimurium(strain_LT2) MAQVINTNSLSLLTQNNLNKSQSALGTAIERLSSGLRINSAKDDAAGQAIANRFTANIKG
G0AIL6_C.fungivorans(strain_Ter3 -MSVINTNILSLVAQNNLTKSASSLSSAIQRLSSGMRINSAKDDAAGQAIANRFTANITG
P04949_E.coli(strain_K12) MAQVINTNSLSLITQNNINKNQSALSSSIERLSSGLRINSAKDDAAGQAIANRFTSNIKG
A0A090GIX0_Mesorhizobium(ORS3359 LSTVQDALGLGASKVDTAYTGMNNVLDTIGKIKTKLLSAV--GQSDANKAKTQTEITALQ
P50612_H.mustelae LGQAISNANDGIGIIQVADKAMDEQLKILDTIKVKATQAAQDGQSLESRKAIQSDIIRLI
P21184_P.aeruginosa LNVATKNANDGISLAQTAEGALQQSTNILQRMRDLSLQSANGSNSDSERTALNGEVKQLQ
P06179_S.typhimurium(strain_LT2) LTQASRNANDGISIAQTTEGALNEINNNLQRVRELAVQSANSTNSQSDLDSIQAEITQRL
G0AIL6_C.fungivorans(strain_Ter3 LTQASRNANDGISLAQTTEGALTEVNNNLQRIRELSVQAASGTNSSSDLQSIQDEVTQRL
P04949_E.coli(strain_K12) LTQAARNANDGISVAQTTEGALSEINNNLQRVRELTVQATTGTNSESDLSSIQDEIKSRL
A0A090GIX0_Mesorhizobium(ORS3359 AQMKSFADAATFSGSNYL----SVTSKQVAAAN---------------------------
P50612_H.mustelae QGLDNIGNTTSYNGQSLLSGQWTNKEFQIGTYSNQSIKVSVGSTTSDKIGQVRINTGAMI
P21184_P.aeruginosa KELDRISNTTTFGGRKLLDGSFGVASFQVGSAANEIISVGIDEMSAES------------
P06179_S.typhimurium(strain_LT2) NEIDRVSGQTQFNGVKVLAQD-NTLTIQVGANDGETIDIDLKQINSQTLGLDTLNVQQKY
G0AIL6_C.fungivorans(strain_Ter3 KEIDRISSQTDFNGVKVLAAN-QTLNIQVGANDGQTIGIDLQKIDSTALKLNTLN-----
P04949_E.coli(strain_K12) DEIDRVSGQTQFNGVNVLAKN-GSMKIQVGANDNQTITIDLKQIDAKTLGLDGFSVKNND
A0A090GIX0_Mesorhizobium(ORS3359 ---------------------DGVQPDAQIVSSFNR------------------------
P50612_H.mustelae TAASEATLTFKQINGGGTSPLEGVKISHSVGTGLGVLAEVINKNSDKTGIRAKASVETTS
P21184_P.aeruginosa --------------------LNGTYFKA----DGGG------------------------
P06179_S.typhimurium(strain_LT2) KVSDTAATVTGYADT--TIALDNSTFKASA-TGLGG----TDQKIDGDLKFDDTTGKYYA
G0AIL6_C.fungivorans(strain_Ter3 --------------------LAGPSGATAAMTPTGK---------DNTG-----------
P04949_E.coli(strain_K12) TVTTSAPVTAFGATTTNNIKLTGITLSTEAATDTGG-----TNPASIEGVYTDNGNDYYA
A0A090GIX0_Mesorhizobium(ORS3359 --------------SSSGAISLGTIDINVESTKLFDSGLSTAVKNQ--------------
P50612_H.mustelae DKEIMSGNLKNL--TINDVNIGNIVDIKKGDAD----GRLVQAINALTSSTGVEAS-TDS
P21184_P.aeruginosa --------------AVTAATASGTVDIAIGITG----GSAVNVKVDMK------GNETAE
P06179_S.typhimurium(strain_LT2) KVTVTGGTGKDGYYEVSVDKTNGEVTLAGGATSPLTGGLPATATEDVKNVQVANADLTEA
G0AIL6_C.fungivorans(strain_Ter3 --------------ASVAVAAGGFTAVNRDAAA----GFTATVADLVT------------
P04949_E.coli(strain_K12) K---ITGGDNDGKYYAVTVANDGTVTMATGATA----NATVTDANTTKATTITSGGTPVQ
A0A090GIX0_Mesorhizobium(ORS3359 -------GILDRKTSIYSTAAD------------------QSAYDAAYA-----------
P50612_H.mustelae KGRLNLRSVDGRGIVLKADASEDNGDGKSAPMAIDAVNGGQSITDGEGAANYGRLSLVRL
P21184_P.aeruginosa QAAAKIAAAVNDANVGIGAFSD--GDTI-SYVS-KA---GKDGSGAITSAVSGVVIA---
P06179_S.typhimurium(strain_LT2) KAALTAAGVTGTASVVKMSYTDNNGKTIDGGLAVKV---GDDYYSATQNKD-GSISI---
G0AIL6_C.fungivorans(strain_Ter3 ---------DARGNTYVKN--------------------GANFYAAT-------------
P04949_E.coli(strain_K12) IDNTAGSATANLGAVSLVKLQDSKGNDTDTYALKDT---NGNLYAADVNETTGAVSV---
A0A090GIX0_Mesorhizobium(ORS3359 ---TAIAGGATDIAANTAGQAAVAGTPVKIDNVS------AFN-----------------
P50612_H.mustelae DARDIVLTSSDKPDENKFSAIGFGDNNVAMATVNLRDVLGKFDASVKSASGANYNAVIAS
P21184_P.aeruginosa ---DTGSTGVGTAAGVTPSATAFA-------------K--TNDTV---------------
P06179_S.typhimurium(strain_LT2) ---NTTKYTADDGTSKT-ALNKLGGADGKTEVVSIGGK--TYAAS--KAEGHNFKAQ---
G0AIL6_C.fungivorans(strain_Ter3 ---VTVADATQGATAATPTATV------------------TYDSS---------------
P04949_E.coli(strain_K12) ---KTITYTDSSGAASSPTAVKLGGDDGKTEVVDIDGK--TYDSA--DLNGGNLQTGLTA
A0A090GIX0_Mesorhizobium(ORS3359 --LDITATGVTDDIITQMVTKIDNVMSQLTDAATVLGAAKSSIDLQKTFTQSLMDSIDRG
P50612_H.mustelae GNSNLGAGVTTLVGAMLVMDIADSARKTLDKIRSDLGSVQGQMVSTVNNISVTQVNVKAA
P21184_P.aeruginosa --AKIDISTAK--GAQSAVLVIDEAIKQIDAQRADLGAVQNRFDNTINNLKNIGENVSAA
P06179_S.typhimurium(strain_LT2) --PDLAEAAAT--TTENPLQKIDAALAQVDTLRSDLGAVQNRFNSAITNLGNTVNNLTSA
G0AIL6_C.fungivorans(strain_Ter3 --KSLTAASGS------PLTAMDSALSQVDSLRSSLGAVQNRFASTIANLGTTVTNLSSS
P04949_E.coli(strain_K12) GGEALTAVANG--KTTDPLKALDDAIASVDKFRSSLGAVQNRLDSAVTNLNNTTTNLSEA
A0A090GIX0_Mesorhizobium(ORS3359 VGQLVDADMNKESTRLQALVQQQLGIQSLSIANSSSQSILQLFKNG
P50612_H.mustelae ESRMREVDFAAESAEFNKYNILAQ----------------------
P21184_P.aeruginosa RGRIEDTDFAAETANLTKNQVLQQAGTAILAQANQLPQSVLSLLR--
P06179_S.typhimurium(strain_LT2) RSRIEDSDYATEVSNMSRAQILQQAGTSVLAQANQVPQNVLSLLR--
G0AIL6_C.fungivorans(strain_Ter3 RSGIQDADYATEVSNMTRSQILQQAGTSVLAKANQSTQSVLTLLQ--
P04949_E.coli(strain_K12) QSRIQDADYATEVSNMSKAQIIQQAGNSVLAKANQVPQQVLSLLQG-

**Figure S3**. Alignment of flagellins used to make two and three species FliC chimeras. Colours indicate flagellin domains (Blue = D0, Orange = D1, Green = D2, Purple = D3).

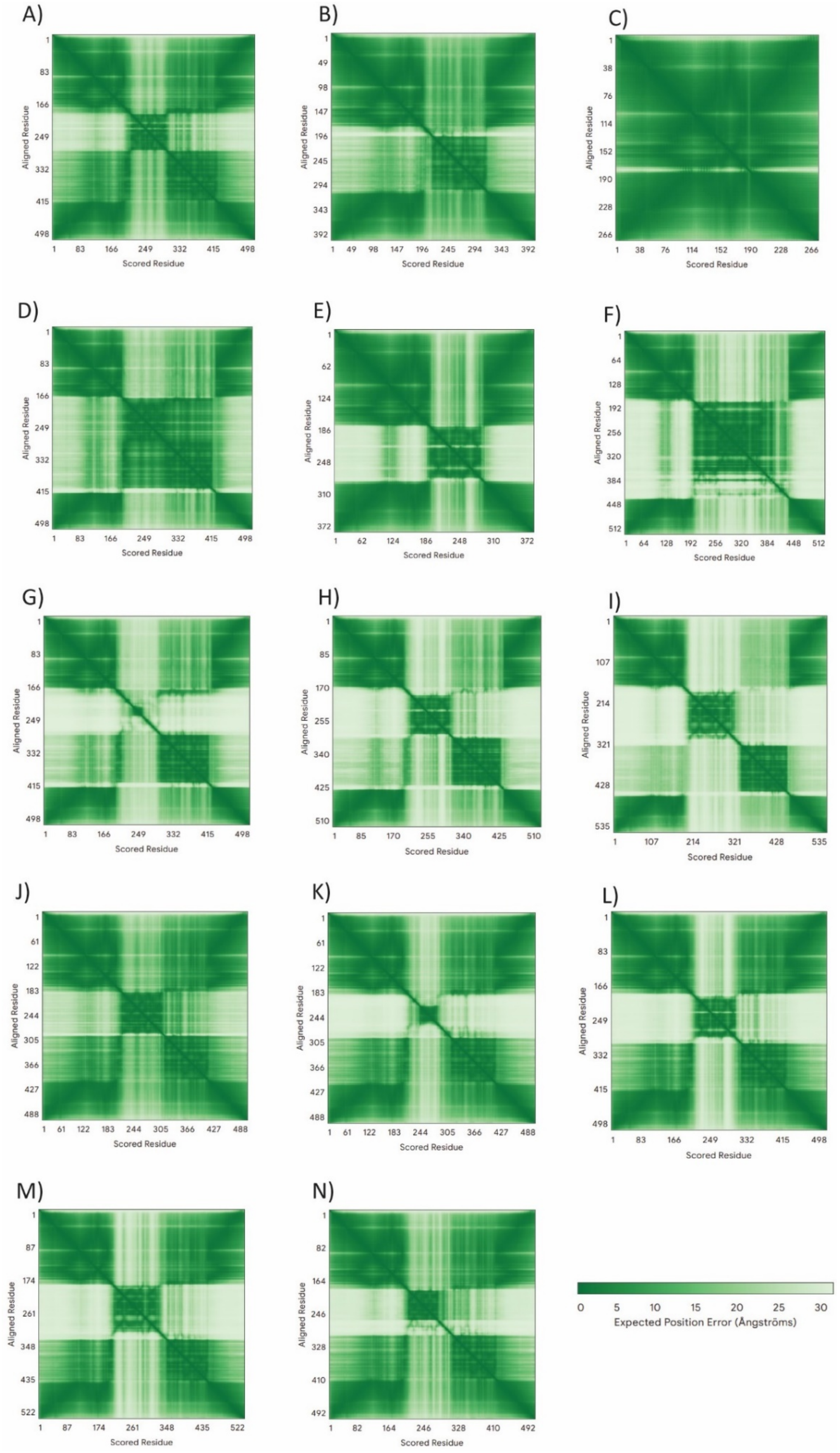

**Figure S4**. Alphafold3 Predicted Aligned Error heatmaps for *E. coli* K-12 FliC and each FliC mutant. Plots of all AlphaFold3 structural predictions for EEEE, EEE, EE, EESS, EEC, EEHH, EESM, EESC,EESP, EEES, EEEM, EEEC, EEEP and EEEH are represented in panels A-N, respectively.

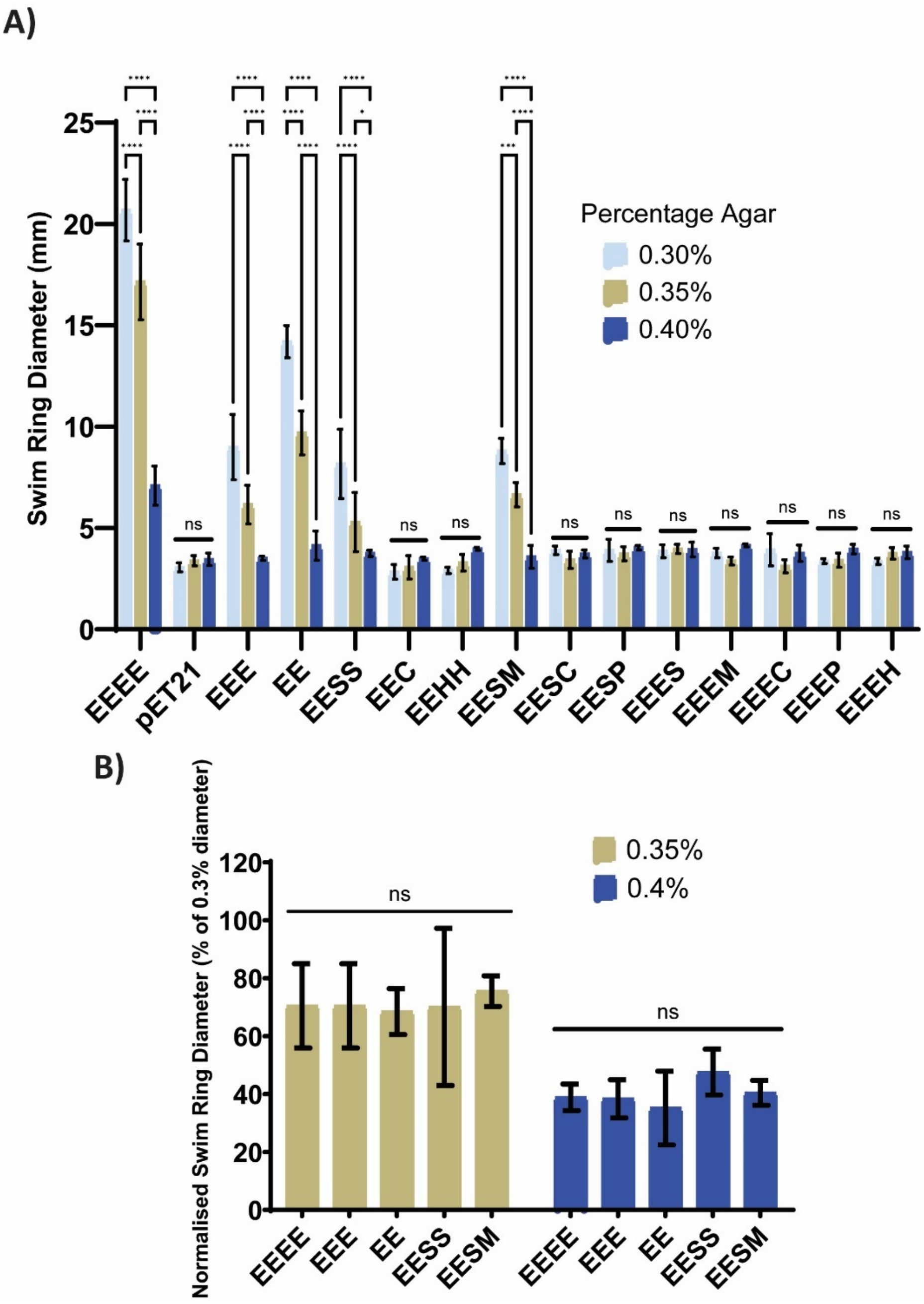

**Figure S5**. A) Swim ring diameters of chimeric fliC expressing *E. coli* strains in 0.3, 0.35, 0.4 and 0.45% agar swim plates. A two-way ANOVA comparing each column to FliC was used to generate the p values (**** = ≤0.0001, *** = ≤0.001and * = ≤ 0.05). B) Normalised percentage swim ring diameters of motile flagellin variants compared to swim rings from 0.3% agar plates. A two-way ANOVA comparing each column to FliC was used to generate the p values (ns = not significant)

**Supplemental Phylogeny S1:** Phylogeny in NEXUS format for phylogeny displayed in Figure 1.

**Supplemental Phylogeny S2:** Phylogeny in NEXUS format for phylogeny displayed in Figure S1.
